# Spindle morphology changes between meiosis and mitosis driven by CK2 regulation of the Ran pathway

**DOI:** 10.1101/2024.07.25.605073

**Authors:** Helena Cantwell, Hieu Nguyen, Arminja Kettenbach, Rebecca Heald

**Author notes:** Corresponding authors: Helena Cantwell and Rebecca Heald.

## Abstract

The transition from meiotic divisions in the oocyte to embryonic mitoses is a critical step in animal development. Despite negligible changes to cell size and shape, following fertilization the small, barrel-shaped meiotic spindle is replaced by a large zygotic spindle that nucleates abundant astral microtubules at spindle poles. To probe underlying mechanisms, we applied a drug screening approach using *Ciona* eggs and found that inhibition of Casein Kinase 2 (CK2) caused a shift from meiotic to mitotic-like spindle morphology with nucleation of robust astral microtubules, an effect reproduced in cytoplasmic extracts prepared from *Xenopus* eggs. In both species, CK2 activity decreased at fertilization. Phosphoproteomic differences between *Xenopus* meiotic and mitotic extracts that also accompanied CK2 inhibition pointed to RanGTP-regulated factors as potential targets. Interfering with RanGTP-driven microtubule formation suppressed astral microtubule growth caused by CK2 inhibition. These data support a model in which CK2 activity attenuation at fertilization leads to activation of RanGTP-regulated microtubule effectors that induce mitotic spindle morphology.

## Introduction

At the meiosis to mitosis transition following fertilization, a rapid shift from the reductional meiotic divisions of the oocyte to the first equational mitotic division of the zygote is required to ensure viable embryonic development. Although cell size and shape remain similar, the cell division machinery is dramatically reconfigured to facilitate this shift in cell division program (Fig 1A). What drives this abrupt change in spindle morphology is an interesting open question.

**Figure 1.**
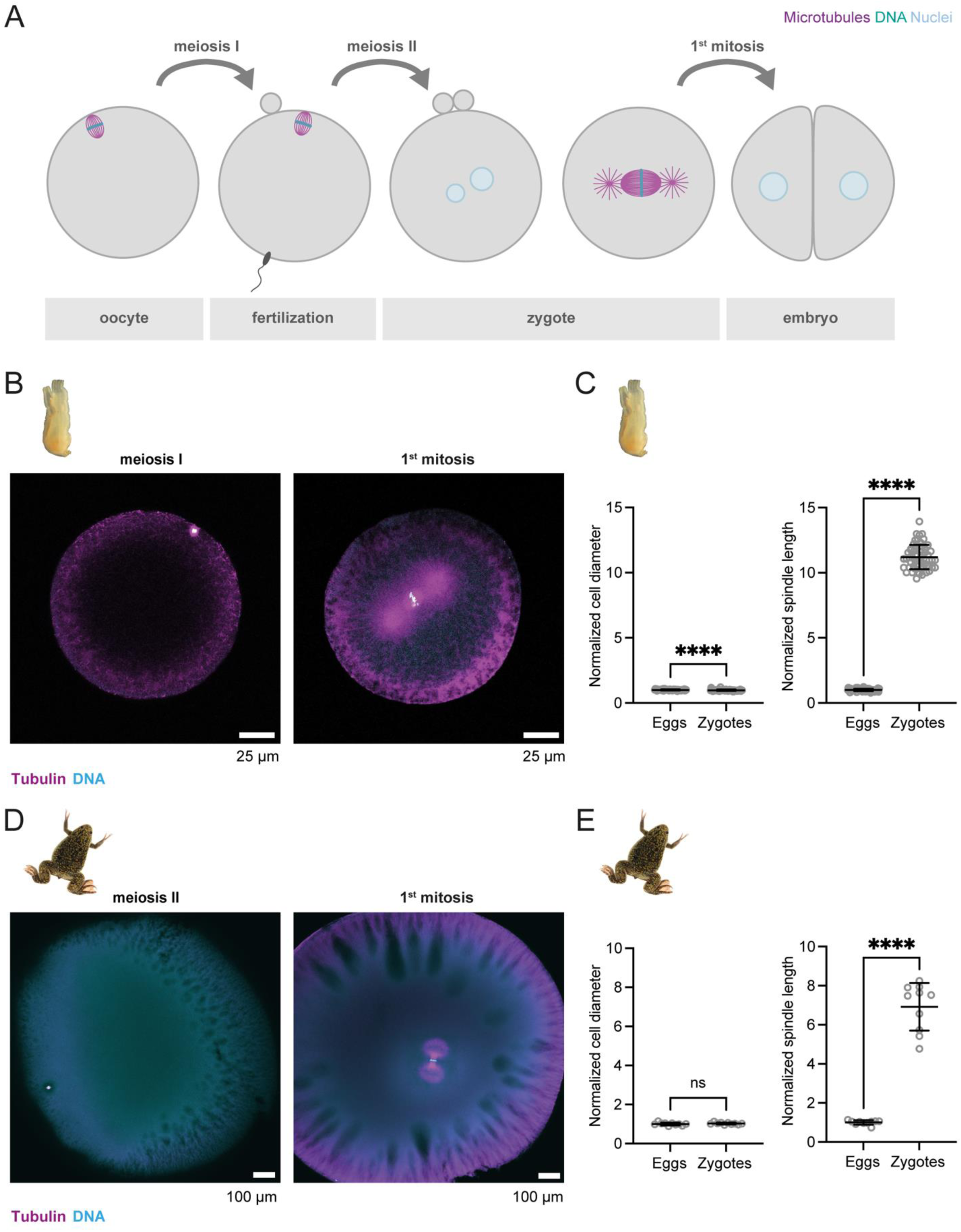
Spindle morphology changes dramatically between meiosis and mitosis. **(A)** Schematic of cellular and spindle morphology changes that accompany the transition from meiotic divisions of the oocyte to the first mitosis of the zygote in early embryogenesis. **(B)** Images of immunofluorescence of a representative *C. robusta* egg in metaphase of meiosis I (left panel) and zygote in metaphase of the first mitosis (right panel). Tubulin (magenta), DNA (cyan). Single z-slice through center of spindle shown. **(C)** Normalized *C. robusta* cell diameter (eggs n=58, zygotes n=58) and metaphase spindle length (eggs n=58, zygotes n=58) in meiosis and mitosis. Measurements are normalized to average egg cell diameter or pole to pole spindle length. Line indicates mean and error bars indicate one standard deviation above and below the mean. **(D)** Images of immunofluorescence of a representative *X. laevis* egg in metaphase of meiosis II (left panel) and zygote in metaphase of the first mitosis (right panel). Tubulin (magenta), DNA (cyan). Single z-slice through center of spindle shown. **(E)** Normalized *X. laevis* cell diameter (eggs n=10, zygotes n=10) and metaphase spindle length (eggs n=10, zygotes n=10) in meiosis and mitosis. Measurements are normalized to average egg cell diameter or spindle length. Statistical significance determined by two-tailed Mann Whitney test (ns = not significant, **** = p < 0.00005). Line indicates mean and error bars indicate one standard deviation above and below the mean.

The spindle is a dynamic, bipolar, microtubule-based structure that in meiosis facilitates the reductional segregation of chromosomes between the oocyte and the much smaller, extruded polar bodies to generate a haploid egg. In mitosis, the spindle faithfully segregates of a copy of each chromosome to two daughter cells. Morphometric differences between meiotic spindles and mitotic spindles of early development are well conserved across metazoan species (Crowder et al., 2015). Meiotic spindles are small, barrel-shaped, anastral across chordates, nematodes, cnidaria and arthropods, and are localized close to the cell periphery. In contrast, mitotic spindles are more centrally positioned in the cell and are larger, scaling with cell size and displaying arrays of astral microtubules emanating from the spindle poles (Crowder et al., 2015).

Although a gradual change in spindle architecture from meiotic to mitotic divisions was reported in mice (Courtois et al., 2012), a more rapid shift in spindle morphology was observed in other metazoan species which, unlike rodents but like humans, inherit a centriole from the sperm upon fertilization (Cavazza et al., 2016; Crowder et al., 2015; Schneider et al., 2021). The sperm centriole subsequently recruits cytoplasmic proteins and duplicates to form two centrosomes, which constitute the major microtubule organizing centers (MTOCs) in mitotic cells. However, the presence of centrioles alone is not sufficient to drive centrosome-mediated astral microtubule nucleation and mitotic spindle morphology. In *Xenopus* egg extracts, spindles formed around sperm chromosomes (with centrosomes) or DNA-coated beads (without centrosomes) both display meiotic anastral morphology, appearing very similar to the meiosis II spindle in unfertilized eggs (Heald et al., 1996). Therefore, activation of microtubule polymerization pathways in the cytoplasm likely drives the change in spindle morphology observed at the meiosis to mitosis transition.

The main pathways responsible for microtubule growth that contribute to spindle assembly include microtubule nucleation and stabilization around chromatin, microtubule nucleation from centrosomes, and branching microtubule nucleation from pre-existing microtubules. One mechanism stimulating microtubule formation around chromosomes is mediated by the chromosome passenger complex (CPC) (Maresca et al., 2009), which consists of INCENP, Borealin, Survivin and Aurora B kinase (Ruchaud et al., 2007; Tseng et al., 2010) and acts to inhibit microtubule depolymerization by MCAK (Sampath et al., 2004) and Stathmin (Gadea and Ruderman, 2006; Kelly et al., 2007), thereby contributing to spindle microtubule stability. A second major pathway for microtubule nucleation and organization around chromosomes is driven by RanGTP, which is generated by its chromatin-bound guanine nucleotide exchange factor RCC1 (Kalab et al., 2002; Oh et al., 2016). A gradient of RanGTP leads to localized release of spindle assembly factors (SAFs) from the transport receptors importin α and importin β that act to sequester them in the cytoplasm by binding to their nuclear localization signal (NLS) sequences (Carazo-Salas et al., 1999; Kalab et al., 2002, 1999; Nachury et al., 2001; Ohba et al., 1999; Wiese et al., 2001; Wilde and Zheng, 1999). A number of importin-regulated SAFs have been identified that play roles in spindle microtubule nucleation and organization (Cavazza and Vernos, 2016) including TPX2 (Gruss et al., 2001; Scrofani et al., 2015), XCTK2 (Ems-McClung et al., 2020, 2004), NuMA (Nachury et al., 2001), and Augmin (Ustinova et al., 2023). Partitioning of importin α between the plasma membrane and the cytoplasm is another mechanism by which NLS-containing cargoes are regulated to promote scaling of spindle size with decreasing cell size during development (Brownlee and Heald, 2019; Rieckhoff et al., 2020).

Another major spindle assembly pathway, active in mitotic cells, is mediated by microtubule nucleation at centrosomes. The centrosome, a pair of centrioles surrounded by pericentriolar material (PCM), concentrates gamma-tubulin ring complexes (γ-TURCs) leading to microtubule nucleation and stabilization (Helmke et al., 2013; Wu and Akhmanova, 2017). Meiotic oocytes lack functional centrosomes but retain centrosomal proteins (Peshkin et al., 2019; Wang et al., 2016), some of which have been shown to function together with PCM components and γ-TURC to nucleate microtubules at acentriolar MTOCs (aMTOCs) (So et al., 2019). In mouse oocytes, a liquid-like domain at spindle poles concentrates regulatory factors promoting spindle assembly (So et al., 2019). Microtubules can also be nucleated on pre-existing microtubules; this branching microtubule nucleation is mediated by the RanGTP-regulated SAFs Augmin and TPX2 and contributes to spindle assembly (Petry et al., 2013; Ustinova et al., 2023; Wu and Akhmanova, 2017). Microtubule plus end binding (+TIP) proteins such as EB3/CLIP-170 and XMAP215/TOG that concentrate tubulin and stimulate microtubule polymerization contribute to this and other nucleation pathways (Kraus et al., 2023; Miesch et al., 2023; Thawani et al., 2018).

How these microtubule nucleation and stabilization pathways are integrated to facilitate formation of the specialized meiotic spindle and, just one cell cycle later, the mitotic spindle of the zygote, remains mysterious. In a study of spindle morphology across a wide range of species, the most dramatic shift in morphology observed at the meiosis to mitosis transition was in the sea squirt *Ciona robusta* (Crowder et al., 2015), a simple chordate species that is the closest living relative of vertebrates (Christiaen et al., 2009a). Here we exploit this robust change in *Ciona* spindle morphology to screen for small molecule inhibitors that perturb it. We then take advantage of the biochemical tractability of *Xenopus laevis* egg and embryo extracts that are capable of recapitulating meiotic and mitotic spindle assembly (Maresca and Heald, 2006; Wilbur and Heald, 2013) to probe candidate pathways and mechanisms. We propose that a decrease in Casein Kinase 2 (CK2) activity following fertilization leads to activation of the RanGTP pathway of microtubule polymerization at spindle poles, thereby contributing to astral microtubule growth and mitotic spindle morphology.

## Results

### Spindle morphology changes dramatically between meiosis and mitosis in both Ciona robusta and Xenopus laevis

To characterize the spindle morphology changes that accompany the meiosis to mitosis transition in chordates (Fig 1A), we visualized microtubules and DNA by immunofluorescence in fixed eggs and zygotes of *Ciona robusta* (*C. robusta)* and *Xenopus laevis (X. laevis). C. robusta* eggs and sperm were isolated from hermaphrodite adults and *in vitro* fertilization was carried out to generate populations of synchronously dividing embryos. *C. robusta* eggs are arrested in metaphase of meiosis I and display a small, barrel-shaped anastral spindle, approximately 10.5 µm in length, close to the cell periphery (Fig 1B, S1A). Though overall cell size and shape of the zygote is similar to that of the egg (Fig 1B, 1C, S1A), the first mitotic spindle in metaphase is approximately 11-fold longer than its meiotic counterpart, more centrally positioned in the cell, and displays large arrays of astral microtubules emanating from the spindle poles (Fig 1B, 1C, S1A); these microtubule arrays extend towards the cell cortex in anaphase (Fig S1B). *X. laevis in vitro* fertilization was carried out to generate synchronously dividing populations of zygotes. *X. laevis* eggs are much larger (∼1.4 mm diameter) than those of *C. robusta* (∼150 µm diameter) (Fig S1A, S1C), and are arrested in metaphase of meiosis II. Like those of *C. robusta, X. laevis* egg meiotic spindles are small (∼26 µm in length), barrel-shaped, anastral and localized in close proximity to the egg periphery whereas zygotic mitotic spindles are approximately 6.9-fold longer than their meiotic counterparts, have astral microtubules emanating from the poles and are more centrally positioned in the cell (Fig 1D, 1E, S1C, S1D).

### CK2 inhibition stimulates astral microtubule assembly and CK2 activity decreases at fertilization in both *Ciona* and *Xenopus*

To identify the upstream pathways underlying the conserved and abrupt change in spindle morphology that occurs following fertilization, we carried out a limited drug screen to identify treatments that perturb meiotic spindle morphology in *C. robusta* eggs. We focused on targets previously reported or hypothesized to modulate known spindle assembly pathways, and then examined spindle morphology by immunofluorescence. In contrast to treatment with the other drugs tested (Fig S2A), treatment with Silmitasertib (CK2i), an inhibitor of Casein Kinase II (CK2), led to a dramatic change in meiotic spindle morphology, pushing it towards a more mitotic-like architecture (Fig 2A). A 1 hour treatment with CK2i caused the majority of oocytes (78%) to form spindles with dramatic arrays of microtubules emanating from the poles and ectopic sites of microtubule nucleation in close proximity to the spindle, with a further 20% displaying a similar but milder phenotype (Fig 2B).

**Figure 2.**
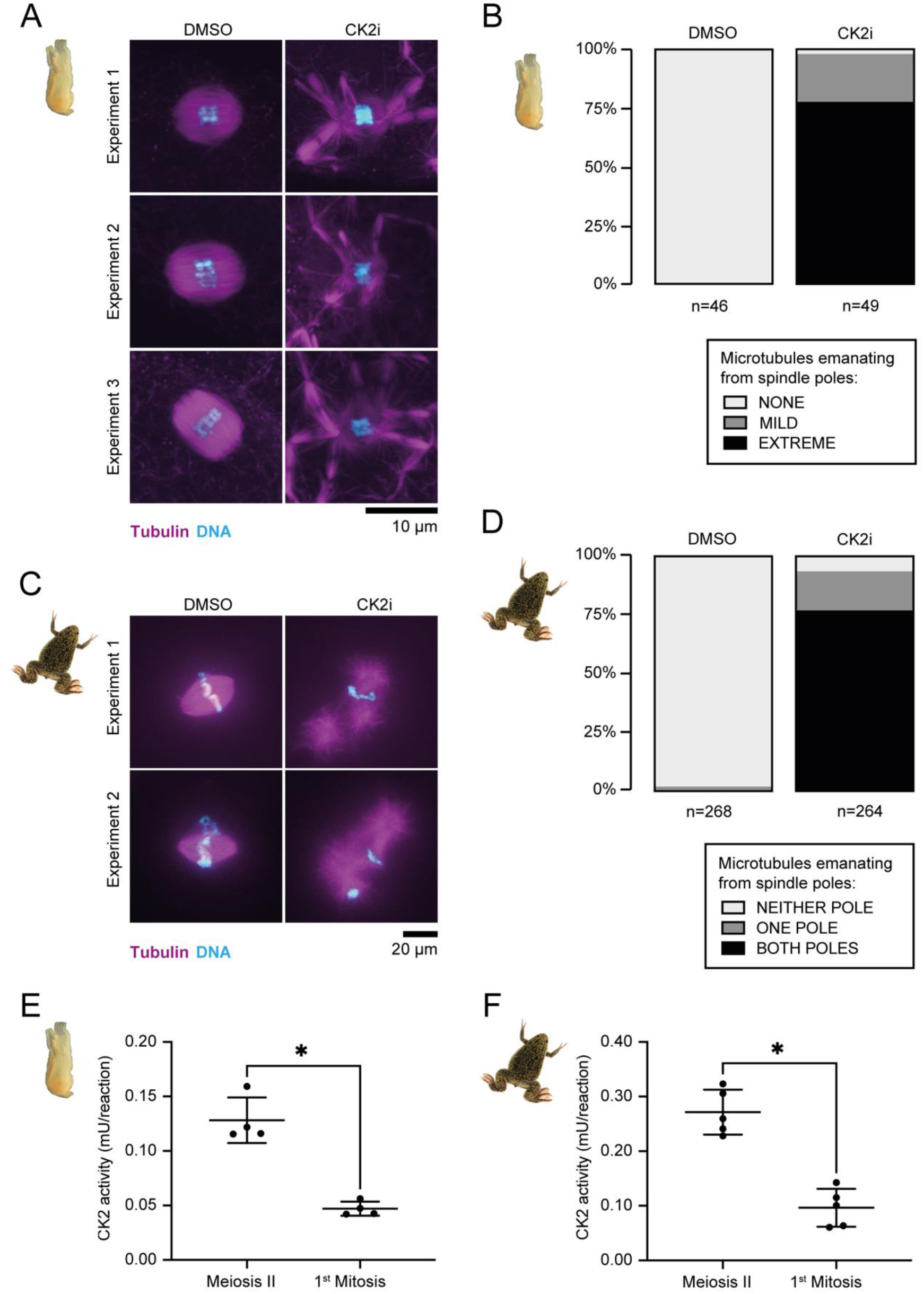
Inhibition of CK2 alters meiotic spindle morphology and CK2 activity decreases between meiosis and mitosis. **(A)** Images of immunofluorescence of representative *C. robusta* oocytes treated for 1 hour with DMSO or 250 µM Silmitasertib (CK2i) as indicated for 3 independent experimental replicates. Maximum intensity projections shown. Tubulin (magenta), DNA (cyan). **(B)** % of *C. robusta* oocytes with spindles displaying no (NONE), limited (MILD) or many (EXTREME) microtubules emanating from the poles following treatment for 1 hour with DMSO or 250 µM CK2i as indicated. Results from 5 independent experiments pooled (n=46 oocytes (DMSO), n=49 oocytes (CK2i)). **(C**) Images of immunofluorescence of representative spindles formed in metaphasic CSF *X. laevis* egg extract in the presence of 50 µM CK2i or DMSO as indicated for two independent experimental replicates. Maximum intensity projections shown. Tubulin (magenta), DNA (cyan). **(D)** Percentage of bipolar spindles displaying microtubules emanating from both poles, one pole or neither pole in metaphasic CSF *X. laevis* egg extract in the presence of 50 µM CK2i or DMSO as indicated. Data from four independent extract experiments pooled (n=264 spindles (CK2i), n=268 spindles (DMSO)); n≥40 spindles per condition for each extract. **(E-F)** CK2 activity (mU/reaction) of cell lysates of meiosis II metaphase and first mitotic metaphase *C. robusta* (E) (n=4 lysates) and *X. laevis* (F) (n=5 lysates) eggs and embryos respectively. Statistical significance determined by two-tailed Mann Whitney test (**** = p < 0.05). Line indicates mean and error bars indicate one standard deviation above and below the mean.

We next assessed the effects of CK2 inhibition on meiotic spindle morphology in *Xenopus* egg extracts (Maresca and Heald, 2006) (Fig 2C). In contrast to the anastral, barrel-shaped spindles formed around *Xenopus* sperm chromosomes in control reactions, CK2i treatment shifted spindle architecture towards a more mitotic-like morphology, with microtubules emanating from the poles (Fig 2C and Fig 2D) and an increased frequency of ectopic microtubule asters in proximity to the spindle (Fig S2B). Therefore, meiotic *Xenopus* cytoplasmic extracts recapitulate the effects of CK2 inhibition observed in *C. robusta* oocytes, suggesting that a potential role for CK2 in regulating spindle microtubules is conserved across chordates.

If attenuation of CK2 activity plays a physiological role in the alteration of spindle morphology observed following fertilization, then its kinase activity is predicted to decrease between metaphase of the second meiotic division and metaphase of the first mitotic division. Utilizing a commercial CK2 kinase activity assay kit with a model substrate we measured CK2 activity in cell lysates of eggs in metaphase of meiosis II and zygotes in metaphase of the first mitosis of both *C. robusta* and *X. laevis.* In both species, CK2 activity decreased between metaphase of the final meiotic and initial mitotic division (Fig 2E and 2F). The observations that CK2 activity decreases at fertilization, and that its inhibition in meiosis leads to mitotic-like spindle morphology, made CK2 a strong candidate to be an upstream regulator of spindle architecture during development.

### Minor CK2-regulated changes to the metaphase proteome accompany the meiosis to mitosis transition

CK2 has been reported to phosphorylate hundreds of different substrates implicated in many signaling pathways and biological processes (Borgo et al., 2021; Meggio and Pinna, 2003). We reasoned that either direct substrates of CK2 or proteins in downstream pathways could regulate spindle morphology. To identify potential candidates at the level of the proteome, we first cataloged changes in protein abundance between meiosis and mitosis and then compared them to those observed upon CK2 inhibition.

To assess overall changes in protein composition between meiosis and mitosis we carried out spindle assembly reactions in metaphase-arrested meiotic egg extracts or in metaphase-arrested mitotic embryo extracts prepared from stage 8 embryos (Wilbur and Heald, 2013) and compared their proteomes by tandem mass tag (TMT) analysis (Fig 3A). To increase coverage, we carried out this analysis twice resulting in two datasets (dataset 1 and dataset 2), each containing three biological replicates.

**Figure 3.**
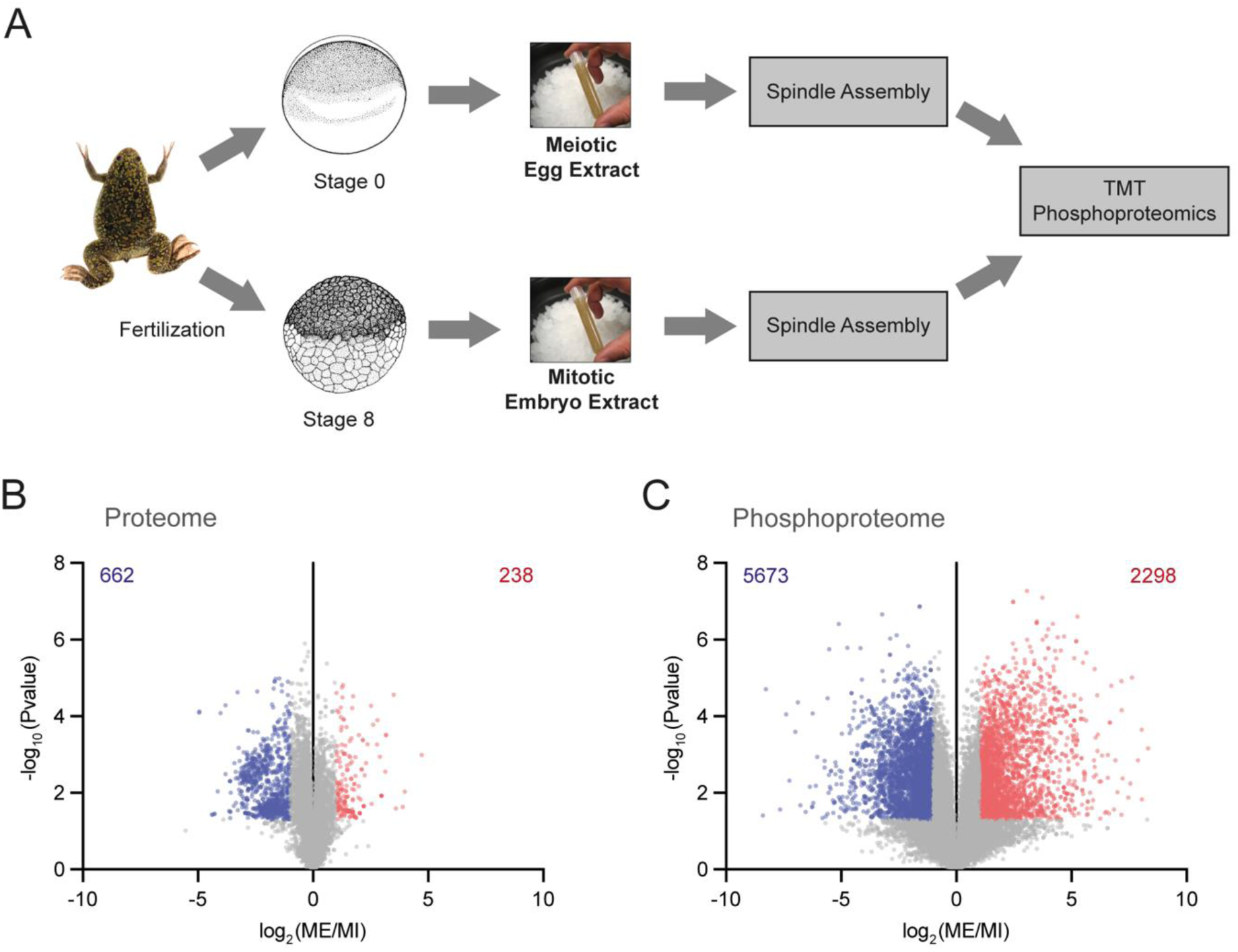
Proteomic and phosphoproteomic changes between meiosis and mitosis. **(A)** Schematic of phosphoproteomics experimental set up. *Xenopus* illustrations © Natalya Zahn (2022) (Fisher et al., 2023; Zahn et al., 2022) **(B)** Volcano plot of log_2_ ratio (protein abundance) versus -log_10_ p value for individual proteins in Meiosis compared to Mitosis (ME/MI). Red points indicate significantly enriched (log_2_ ratio >1 and p value <0.05) proteins (total number indicated in red). Blue points indicate significantly depleted (log_2_ ratio <-1 and p value <0.05) proteins (total number indicated in blue) **(C)** Volcano plot of log_2_ ratio (phosphopeptide abundance) versus -log_10_ p value for individual phosphopeptides in Meiosis compared to Mitosis (ME/MI). Red points indicate significantly enriched (log_2_ ratio >1 and p value <0.05) phosphopeptides (total number indicated in red). Blue points indicate significantly depleted (ratio log_2_ <-1 and p value <0.05) phosphopeptides (total number indicated in blue). Datasets 1 and 2 were combined and visualized together in this figure. Individual datasets are visualized separately in Supplementary Figure S4 (proteome) and Supplementary Figure S5 (phosphoproteome).

At the proteome level (Supplementary Table S1), 13448 proteins were detected (11686 in dataset 1 and 10874 in dataset 2 with 9112 (68%) overlapping between the two datasets). Of these, 238 were enriched in meiosis compared to mitosis (Fig 3B) including Mos, which is degraded upon fertilization (Nishizawa et al., 1993), and Ovochymase, which is secreted upon egg activation (Lindsay et al., 1999). 662 proteins were enriched in mitosis compared to meiosis (Fig 3B). Interestingly, Gene ontology (GO) analysis of proteins overrepresented in meiosis relative to mitosis (Supplementary Table S3) revealed 6 microtubule binding proteins (Supplementary Table S4) including Gamma tubulin complex component 4, a component of the γ-TURC (Oakley et al., 2015). GO categories overrepresented in mitosis relative to meiosis were mostly terms related to cellular membranes and membrane-bound organelles, including the endoplasmic reticulum and Golgi apparatus (Supplementary Table S5).

To determine whether any proteins changing in level between meiosis and mitosis were regulated by CK2, we next compared the proteome of meiotic and mitotic cytoplasmic extracts upon addition of a DMSO solvent control or CK2i at a concentration confirmed by kinase assay to reduce CK2 activity (Fig S3) but that did not perturb the metaphase arrest of the extract as determined by fluorescence microscopy (Supplementary Table S1). Relatively minor changes were seen in the total proteome upon drug treatment (Fig 4A and 4B), with 59 candidate proteins changing in level between meiosis and mitosis and in the same direction upon CK2 inhibition (Fig 4C and Fig 4D). However, none of these proteins possess known roles in microtubule binding or cell division and the only statistically enriched GO category was T cell receptor signaling.

**Figure 4.**
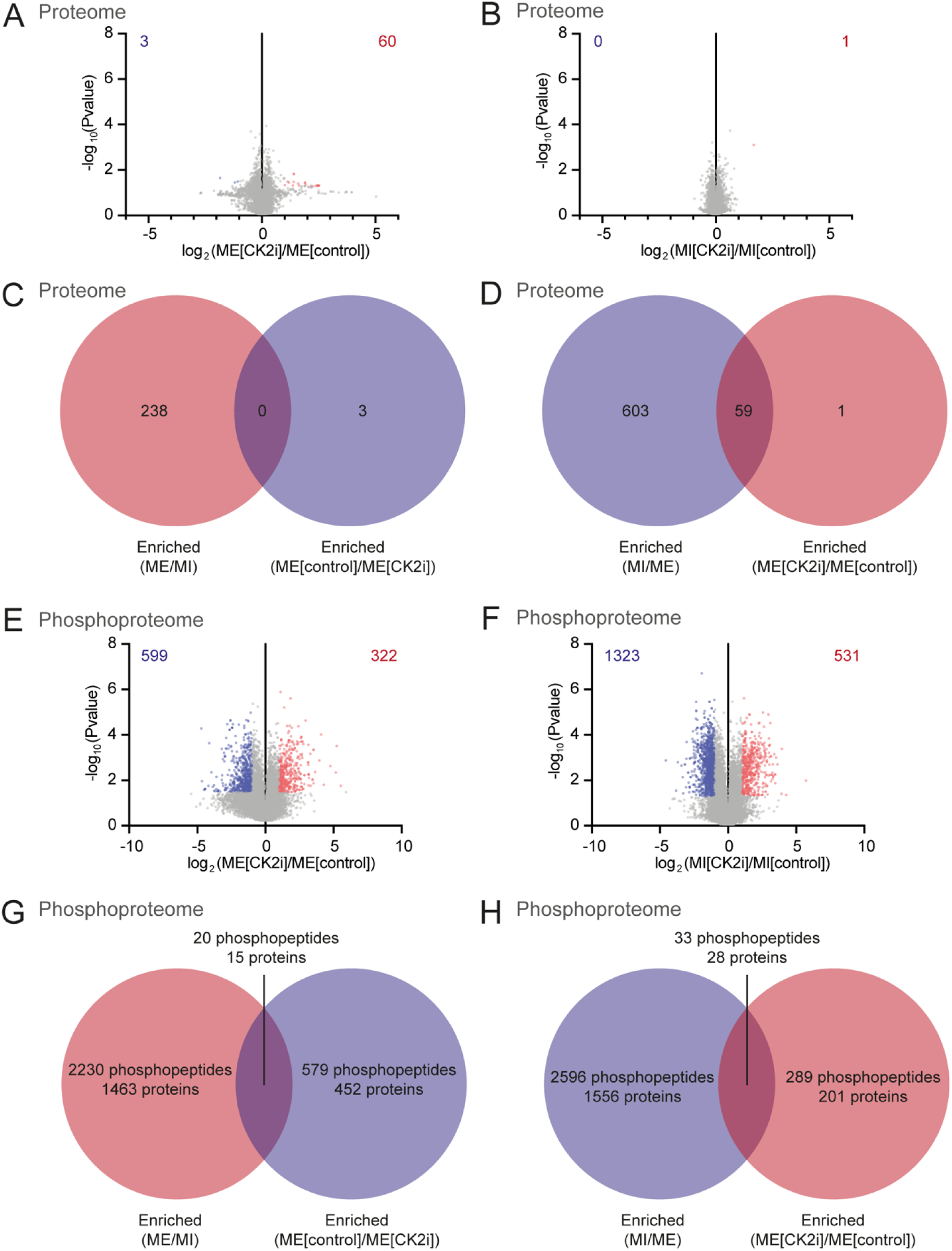
Proteomic and phosphoproteomic changes upon CK2 inhibition. **(A-B)** Volcano plots of log_2_ ratio (protein abundance) versus -log_10_ p value for individual proteins in Meiosis in the presence of CK2 inhibitor (CK2i) compared to Meiosis in the absence of CK2i (A) and Mitosis in the presence of CK2i compared to Mitosis in the absence of CK2i (B) respectively. Red points indicate significantly enriched (log_2_ ratio >1 and p value <0.05) proteins (total number indicated in red). Blue points indicate significantly depleted (log_2_ ratio <-1 and p value <0.05) proteins (total number indicated in blue) **(C)** Venn diagram indicating number of unique proteins in the categories enriched in Meiosis relative to Mitosis and enriched in Meiosis in the absence of CK2i compared to Meiosis in the presence of CK2i and in the overlap between these two categories **(D)** Venn diagram indicating number of unique proteins in the categories enriched in Mitosis relative to Meiosis and enriched in Meiosis in the presence of CK2i compared to Meiosis in the absence of CK2i and in the overlap between these two categories **(E-F)** Volcano plots of log_2_ ratio (phosphopeptide abundance) versus -log_10_ p value for individual phosphopeptides in Meiosis in the presence of CK2i compared to Meiosis in the absence of CK2i (E) and Mitosis in the presence of CK2i compared to Mitosis in the absence of CK2i (F) respectively. Red points indicate significantly enriched (log_2_ ratio >1 and p value <0.05) phosphopeptides (total number indicated in red). Blue points indicate significantly depleted (log_2_ ratio <-1 and p value <0.05) phosphopeptides (total number indicated in blue) **(G)** Venn diagram indicating number of unique phosphopeptides and proteins containing phosphopeptides in the categories enriched in Meiosis relative to Mitosis and enriched in Meiosis in the absence of CK2i compared to Meiosis in the presence of CK2i and in the overlap between these two categories that do not exhibit protein level changes in the same directions **(H)** Venn diagram indicating number of unique phosphopepetides and proteins containing phosphopeptides in the categories enriched in Mitosis relative to Meiosis and enriched in Meiosis in the presence of CK2i compared to Meiosis in the absence of CK2i and in the overlap between these two categories that do not exhibit protein level changes in the same directions. CK2i was added at a concentration of 25 µM in meiotic extract and 10 µM in mitotic extract. Datasets 1 and 2 were combined and visualized together in this figure. Individual datasets are visualized separately in Supplementary Figure S4 (proteome) and Supplementary Figure S5 (phosphoproteome).

Altogether, this analysis reveals some interesting differences between meiotic and mitotic proteomes that may contribute to the change in cell division program. However, since changes in protein levels upon CK2 inhibition were minor, such protein abundance regulation is unlikely to directly underlie CK2 activity-mediated differences between meiotic and mitotic spindle morphology.

### Phosphoproteomics in CK2-inhibited *Xenopus* cytoplasmic extracts identifies candidate phosphoregulated proteins and downstream pathways

Given the rapid shift in spindle microtubule organization at the meiosis to mitosis transition, we reasoned that changes in posttranslational modifications, particularly phosphorylation, were likely to be driving the transition (Peuchen et al., 2017). We assessed the global phosphoproteome and detected 37339 phosphopeptides (Supplementary Table S2). Of these phosphopeptides, 2298 were enriched in meiotic and 5673 in mitotic extracts, corresponding to a total of 1478 and 1584 proteins, respectively (Fig 3C). Whereas GO analysis did not reveal specific categories of proteins with meiosis-enriched phosphosites, 89 spindle-localized proteins were identified with phosphosites enriched in mitosis (Supplementary Tables S6 and S7), including TPX2, HAUS augmin like complex components HAUS5, HAUS6 and HAUS8, TACC3, and kinases Polo 1 kinase and Cyclin dependent kinase 1. We next assessed the effects of CK2 inhibition on the phosphoproteome (Figs 4E and 4F). 53 phosphopeptides from 43 proteins changed in level in the same direction both between meiotic and mitotic extracts and upon CK2i addition to meiotic extract (Fig 4G and Fig 4H). Phase normalization (see Methods) to account for higher total phosphorylation levels in mitotic extract identified a further 35 candidate proteins that change at the phosphopeptide but not the protein level making them candidates for phosphoregulated proteins that drive the change in spindle morphology.

Of the 78 candidate proteins identified by this approach, a total of 38 were previously reported to be associated with the vertebrate spindle or RanGTP-stabilized microtubules by proteomic analyses (Rao et al., 2016; Rosas-Salvans et al., 2018) or by localization to spindles, centrosomes or kinetochores (Huang et al., 2015) (Supplementary Table S8). Network analysis using the STRING database (Szklarczyk et al., 2019) revealed functional clusters of these phosphoregulated proteins with reported roles in DNA replication, nucleocytoplasmic transport, splicing, translation, microtubule binding, chromatin modification, transcription regulation, chromosome condensation, DNA repair and tight junction assembly (Figure 5). Recent work has demonstrated noncanonical roles in cell cycle control for many proteins, notably nucleocytoplasmic transport proteins in spindle assembly (Yang et al., 2023), and our candidates included two proteins reported to be Ran-regulated: nucleoporin NUP214 (Askjaer et al., 1999; Hutten and Kehlenbach, 2006) and cell cycle regulator Geminin (Blow, 2003; Hodgson et al., 2002). Based on literature searches, we hypothesize that a further five candidate CK2-phosphoregulated proteins are likely also Ran-regulated: nucleocytoplasmic transport proteins NUP35 and AKIRIN2 (De Almeida et al., 2021), transcription regulator CTR9 homolog (Kimura et al., 2017; Taltynov et al., 2013), DNA helicase MCM4 (Yamaguchi and Newport, 2003) and telomere-associated protein RIF1 (Sukackaite et al., 2017). Thus, a subset of candidate proteins downstream of CK2 are regulated by RanGTP.

**Figure 5.**
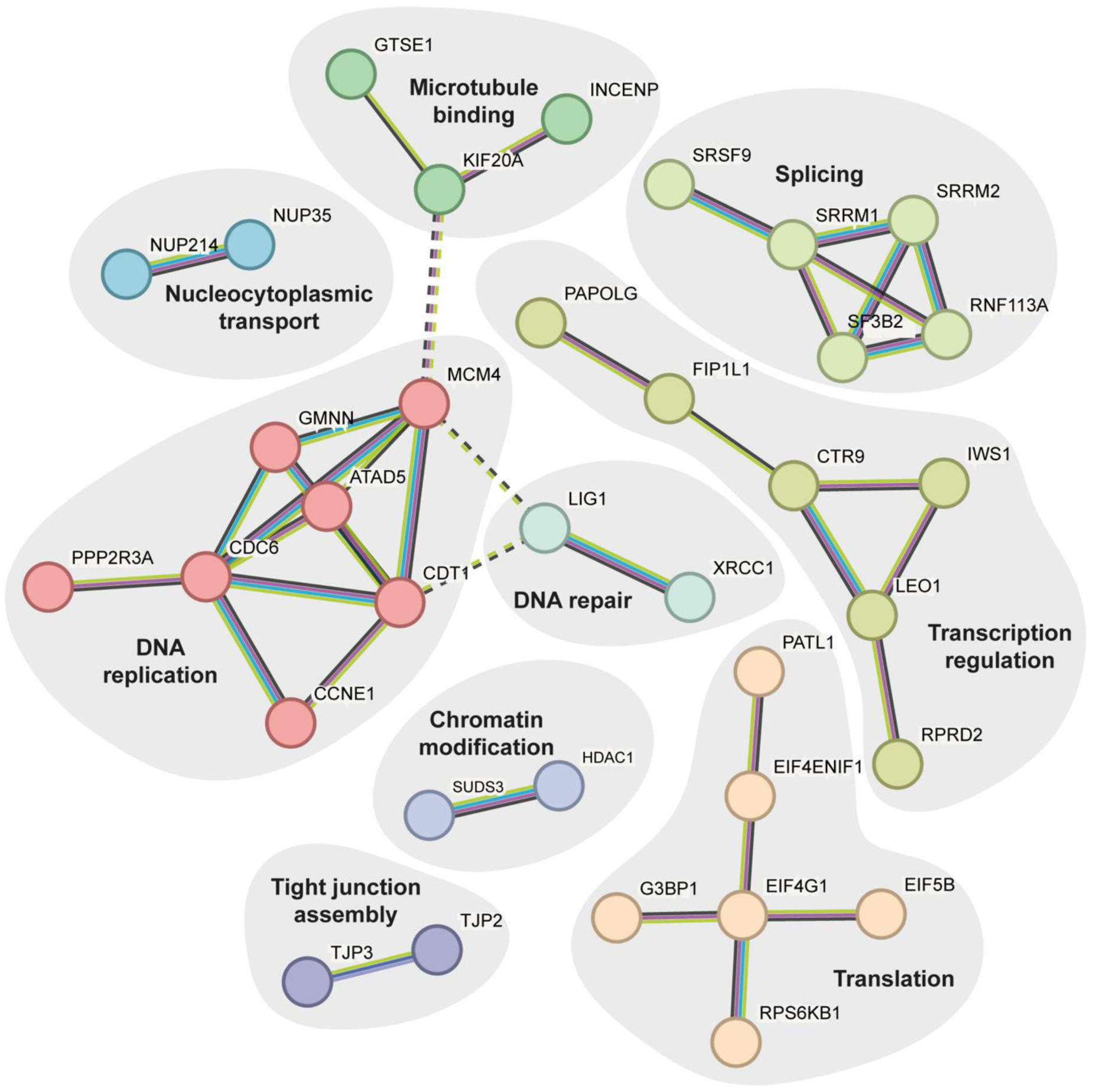
Network analysis of candidate proteins reveals functional clusters. STRING network analysis of candidate proteins carried out using full STRING network annotations with high confidence (interaction score ≥ 0.7). Edge color indicates data source with known interactions shown in cyan (from curated databases) and magenta (experimentally determined), predicted interactions shown in emerald green (gene neighborhood), red (gene fusions) and royal blue (gene co-occurrence), and lime green (textmining), black (co-expression) and lilac (protein homology). K-means clustering with default parameters was used to define functional clusters. Dashed lines represent edges between clusters.

Overall, our phosphoproteomic analysis reveals numerous interesting differences between phosphorylation patterns in meiosis and mitosis, and points to Ran-regulated proteins as potential targets of CK2 regulation that differ between these two distinct cell division programs.

### Inhibition of the RanGTP spindle assembly pathway suppresses effects of CK2 inhibition

To test whether the effect of CK2 inhibition on spindle morphology was mediated by the RanGTP pathway, we used the drug importazole that inhibits the interaction between RanGTP and importin-β (Soderholm et al., 2011). Strikingly, treatment of *C. robusta* oocytes with importazole suppressed the effects of CK2 inhibition on spindle morphology (Fig 6A). Importazole treatment reduced the frequency of oocytes with spindles showing extreme defects with dramatic arrays of microtubules emanating from the poles upon CK2i treatment from 70% to 3%; when treated with both drugs 51% of spindles displayed no microtubules emanating from the poles or ectopic sites of microtubule nucleation in close proximity to the spindle and 46% displayed mild defects, with a small number of microtubule bundles emanating from the poles (Fig 6B).

**Figure 6.**
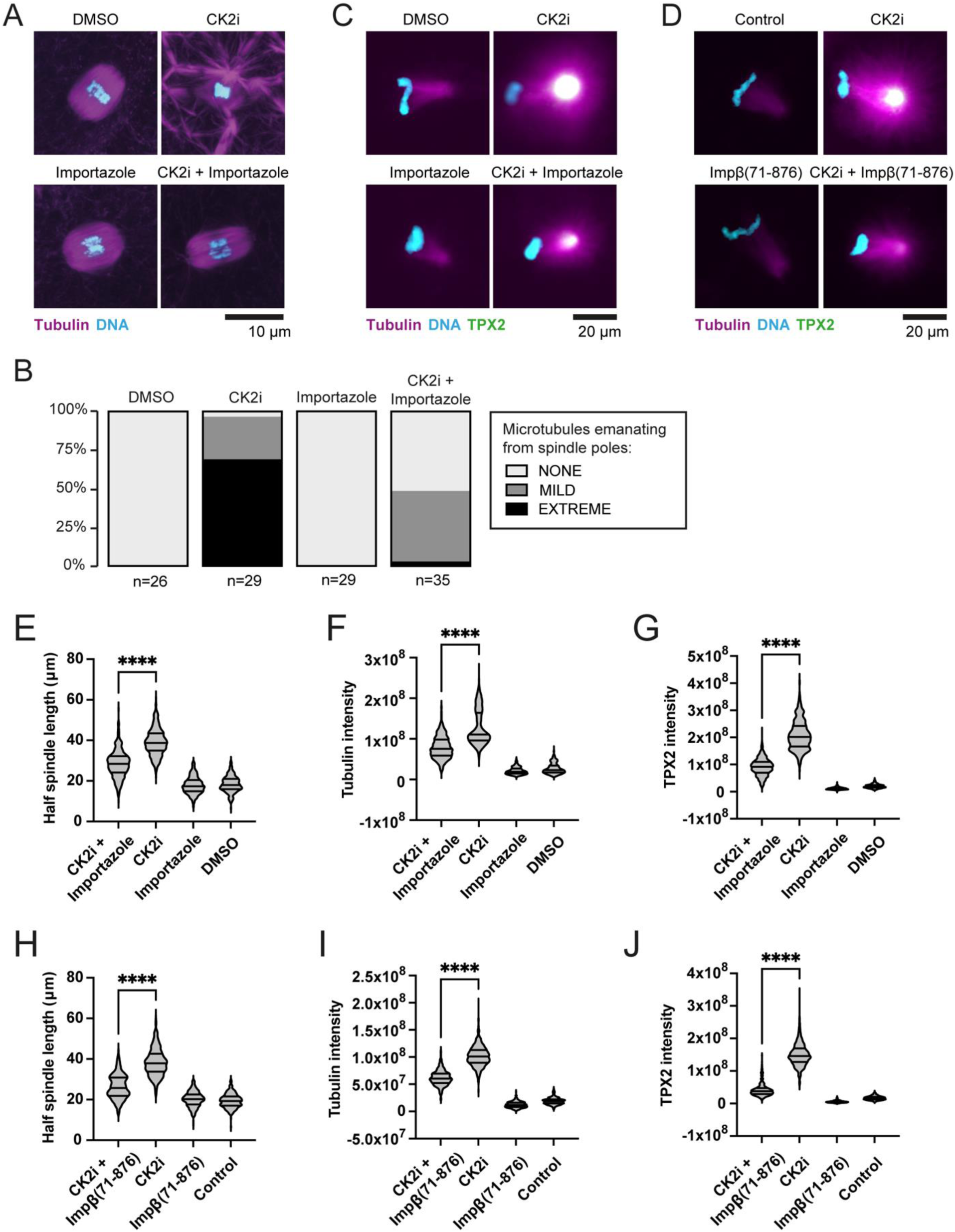
Inhibition of RanGTP-mediated spindle assembly pathway suppresses effects of CK2 inhibition in *C. robusta* eggs and *X. laevis* egg extracts. **(A)** Images of immunofluorescence of representative *C. robusta* oocytes treated for 1 hour with DMSO, 250 µM CK2i, 250 µM importazole or 250 µM CK2i and 250 µM importazole as indicated. Maximum intensity projections shown. Tubulin (magenta), DNA (cyan). **(B)** % of *C. robusta* oocytes with spindles displaying no (NONE), limited (MILD) or many (EXTREME) microtubules emanating from the poles following treatment for 1 hour with DMSO, 250 µM CK2i, 250 µM importazole or 250 µM CK2i and 250 µM importazole as indicated. Results from three independent experiments pooled (n=26 oocytes (DMSO), n=29 oocytes (CK2i), n=29 oocytes (importazole), n=35 oocytes (CK2i + importazole)). Data shown for DMSO and CK2i is a subset of that shown in Fig 2B. **(C)** Representative images of half spindles from spindle assembly reactions carried out in the presence of DMSO, 50 µM CK2i, 300 µM importazole or 50 µM CK2i and 300 µM importazole as indicated. β tubulin (magenta), DNA (cyan), TPX2 (green). **(D)** Representative images of half spindles from spindle assembly reactions carried out in the presence of DMSO and XB (Control), 50 µM CK2i and XB (CK2i), 2.5 µM importin-β(71-876) and DMSO (Impβ(71-876)) or 50 µM CK2i and 2.5 µM importin-β(71-876) (CK2i + Impβ(71-876)) as indicated. β tubulin (magenta), DNA (cyan), TPX2 (green). **(E)** Violin plot of half spindle length of half spindles from experiment described in (C). n≥324 half spindles per condition (from three cytoplasmic extracts (n≥105 per extract for each condition)) **(F-G)** Violin plot of β tubulin (F) or TPX2 (G) intensity of half spindles from experiment described in (C). n≥420 half spindles per condition (from three cytoplasmic extracts (n≥133 per extract for each condition)) **(H)** Violin plot of half spindle length of half spindles from experiment described in (D). n≥313 half spindles per condition (from three cytoplasmic extracts (n≥104 per extract for each condition)) **(I-J)** Violin plot of β tubulin (I) or TPX2 (J) intensity of half spindles from experiment described in (D). n≥329 half spindles per condition (from three cytoplasmic extracts (n≥92 per extract for each condition)). Statistical significance determined by two-tailed Mann Whitney test (**** = p < 0.00005). For violin plots lines indicate median and upper and lower quartiles.

Consistent with a role for RanGTP in mediating the effects of CK2 inhibition, importazole treatment also suppressed the effects of CK2 inhibition on spindle formation when added to *Xenopus* egg extracts. To minimize the effects of redundant mechanisms that drive spindle assembly, we examined microtubule arrays 30 minutes after initiating spindle assembly reactions, when half spindles with a single pole had formed. Half spindles formed in the presence of CK2 inhibitor displayed increased spindle length with large asters at the half spindle poles; importazole treatment suppressed this increase in half spindle length, reducing the size of the aster at the pole (Fig 6C and 6E). The increased intensity of β tubulin at the spindle pole observed upon CK2 inhibition was also suppressed by importazole treatment (Fig 6F). Addition of importin-β(71-876) protein, a truncated mutant that is unable to bind to RanGTP and sequesters cargoes (Nachury et al., 2001), also suppressed the effects of CK2 inhibition on half spindle length and tubulin intensity at the poles (Fig 6D, 6H and 6I).

We hypothesized that the effects of CK2 inhibition on spindle assembly are likely mediated by increased recruitment of importin-β SAF cargoes to spindle poles. To assess this, we monitored the localization of the known SAF TPX2, an importin-β cargo implicated in microtubule nucleation that localizes strongly to the spindle poles in control reactions (Fig S6A). The intensity of both TPX2 and tubulin at the poles was reduced upon addition of importazole or importin-β(71-876) (Fig S6). Both treatments suppressed the increase in TPX2 intensity at the poles observed upon CK2 inhibition (Fig 6C, 6D, 6G, 6J). These results support a model in which a decrease in CK2 kinase activity following fertilization modulates activity of a number of RanGTP-regulated importin cargoes including TPX2, increasing microtubule nucleation and polymerization at spindle poles (Fig 7) and contributing to the difference between meiotic and mitotic spindle morphology.

**Figure 7.**
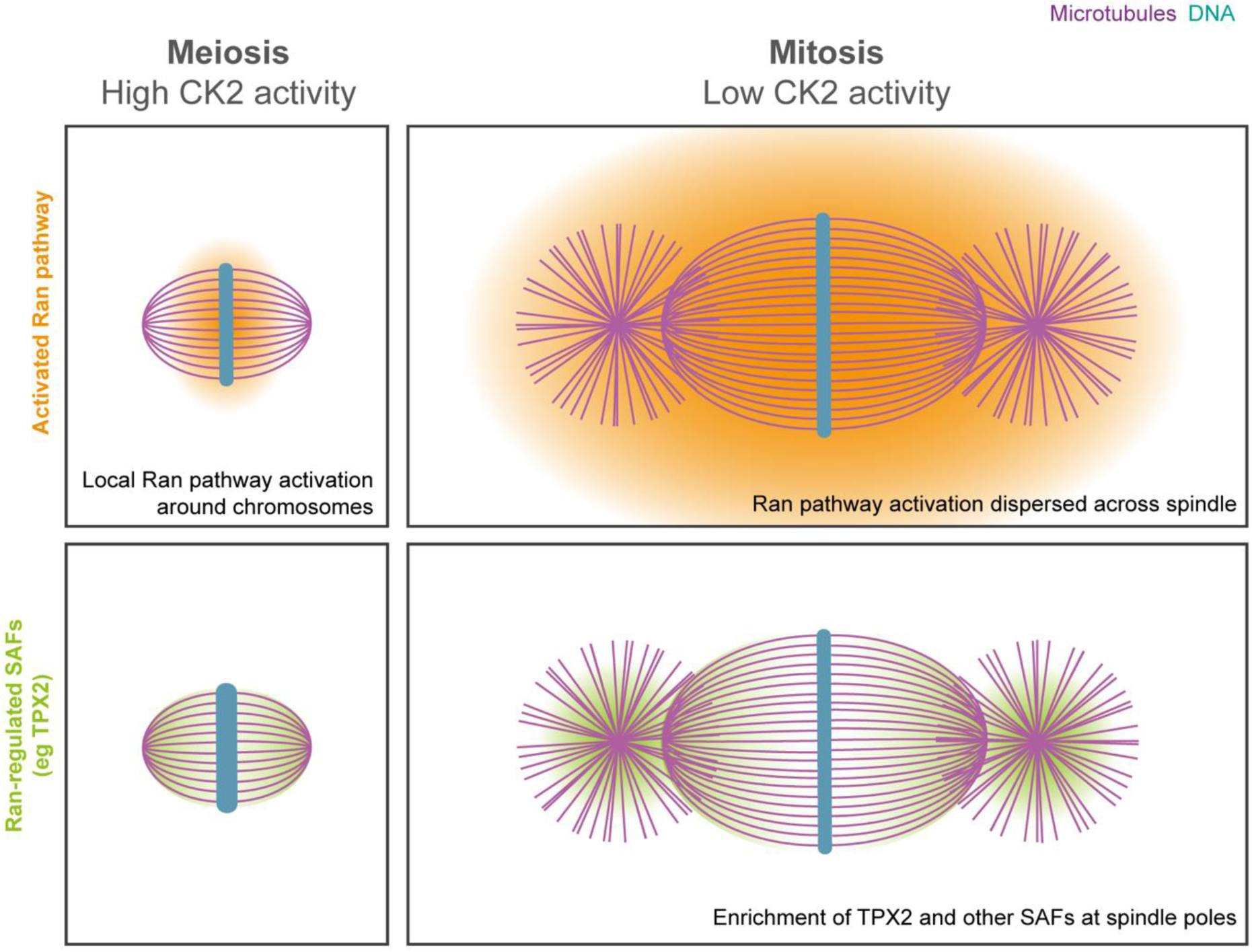
Model of differential regulation of the Ran pathway in meiosis versus mitosis. Schematic illustrating how the Ran pathway of spindle assembly may be altered by the decrease in CK2 activity at fertilization. Microtubules (magenta), DNA (cyan), activated Ran pathway (orange), TPX2 and other Ran-regulated SAFs (green). Where the Ran pathway is highly active, spindle assembly factors (SAFs) are released from importins and drive localized microtubule nucleation and stabilization. In meiosis, Ran pathway activation is limited to a small region surrounding chromosomes. In mitosis, Ran pathway activation extends to a broader area leading to increased localization of TPX2 and other Ran-regulated SAFs, including ELYS and Augmin, at spindle poles, increasing spindle length and driving astral microtubule polymerization at spindle poles.

## Discussion

In this study, we characterized the rapid, dramatic change in spindle morphology between the final meiotic division of the egg and the first mitotic division of the zygote in the sea squirt *C. robusta* and the frog *X. laevis*. Our results provide evidence that the RanGTP pathway is differentially regulated in meiosis and mitosis due, at least in part, to a decrease in CK2 kinase activity at fertilization. In *Ciona*, this drop in activity is likely caused by degradation of regulatory subunit CK2β (Russo et al., 2004), while mechanisms regulating CK2 activity in *Xenopus* remain unclear and do not appear to be at the proteome level.

Inhibition of CK2 resulted in the formation of astral microtubule networks that mimic the effects of adding constitutively active RanGTP to meiotic *Xenopus* egg extract (Kalab et al., 1999). That targets of Ran are regulated by CK2 is supported by phosphoproteomic analysis. In addition, we observed enhanced spindle pole localization of the Ran-regulated SAF TPX2 upon CK2 inhibition. Strikingly, this effect was reversed by inhibition of cargo release from importins by either a drug or recombinant protein, suggesting a global effect on the RanGTP pathway at the meiosis to mitosis transition that stimulates localization of SAFs at spindle poles and astral microtubule formation. One possible explanation is that CK2-mediated phosphorylation inhibits RanGAP, its GTPase activating protein that forms a complex with Ran and RanBP1 and promotes hydrolysis of RanGTP to RanGDP. CK2 phosphorylation of RanGAP1 on serine 358, a site conserved in *Xenopus*, has been shown in humans to increase the efficiency of RanGAP1-Ran-RanBP1 complex formation, which could affect Ran-GTP hydrolysis rate and or the spatial activation of SAFs (Caudron et al., 2005; Gruss and Vernos, 2004; Kalab and Heald, 2008; Takeda et al., 2005). Interestingly, partitioning of importin α between cellular membranes and the cytoplasm, where it binds and inhibits SAFs, is also regulated by CK2-mediated phosphorylation (Brownlee and Heald, 2019; Hachet et al., 2004). Unfortunately, these phosphosites on RanGAP and importin α were not present in our phosphoproteomics dataset, and further work will be needed to investigate their regulation by CK2 at the meiosis to mitosis transition.

The RanGTP pathway has previously been implicated in meiotic spindle assembly in multiple systems (Cavazza and Vernos, 2016; Cesario and McKim, 2011; Drutovic et al., 2020; Dumont et al., 2007; Gruss et al., 2001). Our data indicate that the RanGTP pathway, differentially regulated, is also important for determining mitotic spindle morphology in early embryos. This supports the conclusions of recent work that demonstrated that the RanGTP pathway is essential for mitotic spindle assembly during the very early embryonic cleavage divisions of medaka fish embryos (Kiyomitsu et al., 2024) and work showing that spindle assembly in *X. laevis* early embryo cytoplasmic extracts (Stage 3) was dependent on the RanGTP pathway (Wilbur and Heald, 2013).

The first mitotic division of human embryos is frequently error prone (Cavazza et al., 2021; Currie et al., 2022) contributing to the high rates of aneuploidy in early embryos (Coticchio et al., 2023; Lee and Kiessling, 2017). Very little is known about what regulates the change in spindle morphology at the shift from egg to zygote, a process critical for accurate chromosome segregation in the first mitotic division. Our study implicates CK2 and the RanGTP pathway in regulation of this spindle morphology transition in both frogs and sea squirts suggesting likely conservation of regulatory mechanisms across species. Overexpression of HSET (kinesin-14) in mouse oocytes elongated spindles, making them more mitotic-like (Bennabi et al., 2018) and expression of mutant INCENP in *Drosophila* oocytes led to ectopic aster formation in proximity to the spindle (Colombié et al., 2008) suggesting these factors may also contribute to the spindle morphology change. A recent preprint demonstrated that inhibition of the Mos-MAPK pathway led to a shift in spindle morphology to mitotic-like, with astral microtubules, in starfish oocytes (Avilov et al., 2023 *Preprint*) suggesting this signaling pathway, which is active in meiosis but inactivated by degradation of Mos at fertilization (Dupré et al., 2011), may be part of the upstream control.

Here we have focused on spindle morphology but our global phosphoproteomic approach also provides a dataset in which we can probe changes in the regulation of other cellular processes that accompany the meiosis to mitosis transition. Changes to chromosome condensation (Bomar et al., 2002) and cohesion (Tachibana-Konwalski et al., 2010), cell cycle regulation (Kubiak et al., 2008) and centrosome regulation (Manandhar et al., 2005) accompany the transition from egg to zygote. Interestingly, proteins with phosphosites enriched in mitosis compared to meiosis in our dataset included condensins, NUSAP1 and histones H1.3 and H1.8, and 74 centrosomal proteins suggesting phosphoregulation of chromosome condensation and centrosome regulation between meiosis and mitosis. Future work will explore the molecular basis of these changes and the contribution of phosphoregulation to ensuring they are brought about accurately at the appropriate time in development to ensure embryonic viability.

## Materials and methods

### *Ciona* husbandry, gamete collection and *in vitro* fertilization

*C. robusta* adult animals were obtained from M-Rep and maintained at 16°C in artificial seawater (Instant Ocean). Gametes were obtained by dissection as described previously (Christiaen et al., 2009b). Eggs and sperm from 6-12 animals were mixed for each experiment. For *in vitro* fertilizations, sperm was activated by mixing with basic artificial seawater (10µl 1M TRIS pH9/10 in 6ml seawater) in a gelatin-coated petri dish. Eggs were mixed with sperm for 3 min then dechorionated by mixing with dechorionation solution (0.1 g/ml sodium thioglycolate, 0.01 g/ml protease (*Streptomyces griseus*) (Sigma PP8811), 0.06M NaOH) and then washed three times with artificial seawater and embryos maintained at 16°C.

### *Xenopus* husbandry, gamete collection and *in vitro* fertilization

Mature *X. laevis* were obtained from *Xenopus*1 or the National *Xenopus* Resource (Woods Hole) and maintained and used following standard protocols in the Animal Use Protocol approved by UC Berkeley Animal Care and Use Committee. J-strain (back-crossed) *X. laevis* (National *Xenopus* resource) were used to prepare samples for all proteomic and phosphoproteomic experiments. Testes were dissected from adult males and stored at 4°C in MR (100 mM NaCl, 1.8 mM KCl, 1 mM MgCl_2_, 2 mM CaCl_2_, 5 mM HEPES (pH 7.6)) for less than one week. Females were primed by injection with 100 U of pregnant mare serum gonadotropin (PMSG) (National Hormone and Peptide Program) at least 48h before use and boosted by injection with 500 U of human chorionic gonadotropin (hCG) 16h before use. Females were gently squeezed to deposit eggs into petri dishes. 1/3 of a teste was homogenized using scissors and a plastic pestle in 1.1 ml ddH_2_O in a 1.5 ml tube then added to the petri dish of eggs and swirled to mix and incubated for 5-10 min. Dishes were flooded with 0.1X MMR (100 mM NaCl, 2 mM KCl, 1 mM MgCl_2_, 2 mM CaCl_2_, 5 mM HEPES (pH 7.6), 0.1 mM EDTA) and incubated for 10 min. Jelly coats were removed by incubation in 2% cysteine solution (in ddH_2_O-NaOH (pH 7.8)), then embryos were washed at least four times in 0.1X MMR and incubated at 23°C.

### *Xenopus* egg and embryo extract preparation and spindle assembly reactions

*X. laevis* crude CSF egg extracts were prepared as described previously (Maresca and Heald, 2006). Eggs were packed using a clinical tabletop centrifuge and then crushed by centrifugation for 16 min at 10,200 rpm (16°C) in an HB-6 rotor (Sorvall). Cytoplasm was removed and supplemented with 20 µg/ml Cytochalasin B (Cyto B), 10 µg/ml LPC (leupeptin, pepstatin, chymostatin), 1X energy mix (3.75 mM creatine phosphate, 0.5 mM ATP, 0.05 mM EGTA, 0.5 mM MgCl_2_). Stage 8 embryo extracts were prepared as described previously (Wilbur and Heald, 2013). Embryos maintained for 5.5h post fertilization at 23°C were washed extensively in CSF-XB (5 mM EGTA, 100 mM KCl, 2 mM MgCl_2_, 0.1 mM CaCl_2_, 50 mM sucrose, 10 mM HEPES (pH 7.8)) supplemented with 20 µg/ml Cyto B and packed by centrifugation at 1000 rpm for 1 min then 2000 rpm for 30 secs in a microcentrifuge at 16°C then crushed by centrifugation for 12 min at 10,200 rpm (16°C) in an HB-6 rotor (Sorvall). Cytoplasm was removed and supplemented with 20 µg/ml Cyto B, 10 µg/ml LPC, 1X energy mix, 0.2 mg/ml UbcH10 C114S, 0.05 mg/ml Cyclin B delta 90.

For spindle assembly reactions, purified *X. laevis* sperm nuclei were added to egg and embryo extracts at a concentration of 1000 nuclei/µl. To visualize microtubules extracts were supplemented with 0.3 µM rhodamine-labeled tubulin. For sample preparation for proteomic and phosphoproteomic analysis rhodamine-labeled tubulin was not added but spindle assembly was monitored by microscopy in a parallel test reaction using the same extract and conditions labeled with rhodamine-tubulin.

### Fixation, immunofluorescence and imaging

*C. robusta* eggs and embryos were fixed overnight at 4°C in ‘von Dassow fixative’ (100 mM HEPES (pH 7), 50 mM EGTA (pH 7), 10 mM MgSO_4_, 500 mM dextrose, 2% formaldehyde, 0.2% glutaraldehyde, 0.2% triton-X100) (Crowder et al., 2015) in 5% BSA-coated tubes, washed three times in PBT (PBS, 0.1% triton-X100), once in PBS and incubated in 0.1% NaBH_4_ in PBS. For immunofluorescence, *Ciona* samples were blocked (5% normal goat serum, 0.2% BSA in PBT) overnight at 4°C, incubated in primary antibody (1.25 µg/ml E7 anti-β tubulin (DSHB) in PBT + 0.2% BSA) for 72h at room temperature (RT), washed for 24h in PBT + 0.2% BSA, incubated in secondary antibody (2 µg/ml goat anti-mouse Alexa Fluor 568 (Thermo Fisher Scientific) for 72h at RT, washed for 24h in PBT + 0.2% BSA, incubated in 1 µg/ml hoescht in PBT + 0.2% BSA for 30 min at RT, washed three times in PBT then mounted in vectashield (vector labs) between tape spacers on a glass slide. Samples were imaged using a Zeiss PlanApo 63x (NA 1.4) oil or Zeiss PlanApo 20x (NA 0.8) air objective on a Zeiss LSM 800 confocal microscope with 488 and 568 laser lines. Egg and zygote cell diameter and spindle pole-to-pole length measurements were made using Imaris 9.5.1.

*X. laevis* eggs and embryos were fixed for 1-3h at RT in MAD fixative (40% MeOH, 40% Acetone, 20% DMSO) then stored at -20°C. Samples were gradually rehydrated into 0.5X SSC (75 mM NaCl, 15 mM sodium citrate, pH 7) then bleached under direct light in bleaching solution (2% H_2_O_2_, 5% formamide in 0.5X SSC) for 2-3h and washed twice in PBT. Samples were blocked in 10% normal goat serum + 5% DMSO in PBT overnight at 4°C then incubated at 4°C for 24h in primary antibodies (1.25 µg/ml E7 anti-β tubulin (DSHB) and 2 µg/ml ab1791 anti-H3 (abcam) in PBT + 10% normal goat serum), washed for 24h in PBT, incubated for 24h at 4°C in secondary antibodies (4 µg/ml goat anti-mouse Alexa Fluor 568 (Thermo Fisher Scientific) and 4 µg/ml goat anti-rabbit Alexa Fluor 488 (Thermo Fisher Scientific) in PBT) then washed for 24h in PBT. Samples were then gradually dehydrated into 100% MeOH, stored overnight at -20°C then cleared by incubation in Murray’s clear (2 parts benzyl benzoate, 1 part benzyl alcohol) then mounted on a glass coverslip. Samples were imaged using a Zeiss PlanApo 20x (NA 0.8) air objective on a Zeiss LSM 800 confocal microscope with 488 and 568 laser lines. Egg and zygote cell diameter and spindle length measurements were made using Imaris 9.5.1. Cell diameter was measured across widest part of cell. Spindle length was measured from outer edge of aster to edge of aster, or pole to pole for meiotic spindles with no asters were present, along the long axis of the spindle.

Spindles and half spindles from *X. laevis* spindle assembly reactions were spun down onto coverslips and fixed as described previously (Hannak and Heald, 2006). Briefly, spindles or half spindles were fixed in spindle dilution buffer (30% glycerol, 1 X BRB80 (80 mM PIPES (pH 6.8), 1 mM MgCl_2_, 1 mM EGTA), 0.5% Triton-X100, 3.7% formaldehyde) then spun down onto glass coverslips through a cushion (40% glycerol, 1X BRB80) by centrifugation for 20 min at 5,500 rpm (16°C) in an HS-4 rotor (Sorvall). Coverslips were post-fixed for 5 min in cold 100% methanol then washed in PBS NP40 (1X PBS, 0.1% NP40). For immunofluorescence, coverslips were blocked with PBS-BSA (1X PBS, 3% BSA) for 45 min at RT, incubated at 4°C overnight with primary antibody (1:4000 rabbit-anti-TPX2 or 1.67 µg/ml 3753 rabbit-anti-GTSE1 (gift from Alexander Bird)) in PBS-BSA, washed with PBS-NP40, incubated for 1h at RT with secondary antibody (4 µg/ml goat anti-rabbit Alexa Fluor 488 (Thermo Fisher Scientific) in PBS-BSA) then washed with PBS-NP40. To visualize DNA, coverslips were incubated for in 1 µg/ml hoescht in PBS-NP40 then washed with PBS-NP40. Coverslips were mounted in vectashield (vector labs) on a glass slide and sealed with clear nail polish. Samples were imaged using an Olympus UPlan Fl 40X (NA 0.75) air objective and Hamamatsu ORCA-ER camera on an Olypmus BX51 widefield microscope or a Zeiss Plan-Apochromat 40X (NA 0.95) air objective and Axiocam 712 camera on a Zeiss Axio Observer 7 widefield microscope (using Apotome 3 for representative images). Half spindle length measurements were made using Fiji (ImageJ). Half spindle length was measured as the distance from edge of aster (or pole if no aster was present) to DNA. To quantify fluorescence intensity at half spindle poles total fluorescence intensity in each channel was measured in a square (200 pixels x 200 pixels) centered on each spindle pole and total fluorescence intensity of a background square (200 pixels x 200 pixels) was subtracted from it.

### Drug treatments and protein additions

For limited drug screen, *Ciona* eggs and embryos were bathed in artificial sea water containing the indicated concentration of Silmisatertib (MedChem Express), Importazole (Sigma-Aldrich), Palmostatin (Millipore-Sigma), Monastrol (Abcam), BI2536 (Selleck Chemicals), Okadaic acid (Cayman Chemical Company), ZM447439 (Selleck Chemicals) or Tozasertib (MedChem Express). For drug treatments of extracts, drugs stocks were diluted 1:5 in XB (1 mM MgCl_2_, 0.1 mM CaCl_2_, 100 mM KCl, 50 mM sucrose, 10 mM HEPES (pH 7.7)) and added at the beginning of the spindle assembly reaction. For protein addition to extracts, proteins were in XB and an equal volume of XB was added to the control reaction.

### CK2 activity assay

For *X. laevis* lysates, 10 dejellied eggs or zygotes of the appropriate stage (confirmed by immunofluorescence of parallel sample taken at the same timepoint after fertilization) were snap frozen. Samples were resuspended in 50µl of *Xenopus* lysis buffer (25 mM HEPES (pH 7.2), 10 mM EDTA, 250 mM sucrose, 1% NP40, 1 tablet/10 ml PhosSTOP (EDTA-free) (Roche), 1 tablet/10 ml cOmplete mini protease inhibitor (EDTA-free) (Roche), 0.2 mM phenylmethylsulfonyl fluoride (PMSF), 10 µM cytochalasin D) on ice and lysed by pipetting then thorough vortexing. Yolk removal was carried out as previously described (Van Itallie et al., 2021, *Preprint*); samples were centrifuged at 4,000 g for 4 min (4°C) in a microcentrifuge then lipids were resuspended by gentle flicking and the supernatant was retained. Total protein concentration of samples was determined by Bradford assay (Bio-Rad) and samples were diluted to a concentration of 500 µg/ml. To determine CK2 activity, a CycLex CK2 Kinase Assay Kit (MBL Life Science) was used according to the manufacturer’s recommended protocol.

For *C. robusta* lysates, dechorionated eggs or zygotes of the appropriate stage and from a fertilization with a fertilization efficiency of <79% (confirmed by immunofluorescence of parallel sample taken at the same timepoint after fertilization) were washed in PBS, pelleted by centrifugation at 3000 rpm for 1 min (16°C) in a microcentrifuge and snap frozen. Pellets were resuspended in 100 µl *Ciona* lysis buffer (20 mM TRIS-HCl (pH 7.4), 1 mM EDTA, 1 mM EDTA (pH 8.0), 150 mM NaCl, 0.5% NP40, 1 tablet/10 ml PhosSTOP (EDTA-free) (Roche), 1 tablet/10 ml cOmplete mini protease inhibitor (EDTA-free) (Roche), 0.2 mM PMSF) on ice, incubated for 5 min on ice and then lysed with a plastic pestle for 1 min. Following centrifugation at 13,000 rpm for 5 min (4°C) the supernatant was retained. Total protein concentration of samples was determined by Bradford assay (Bio-Rad) and samples were diluted to a concentration of 500 µg/ml. To determine CK2 activity, a CycLex CK2 Kinase Assay Kit (MBL Life Science) was used according to the manufacturer’s recommended protocol.

### Sample preparation for proteomics

For each of 2 experimental repeats, spindle assembly reactions were carried out in 3 independent meiotic egg extracts and 3 independent mitotic embryo extracts in the presence of 25 µM Silmisatertib or DMSO (meiotic) or 10 µM Silmisatertib or DMSO (mitotic). After 1h incubation at 18°C, once spindles had formed, samples were snap frozen in liquid nitrogen and stored at -80°C. Total protein concentration of the sample was determined by Bradford assay (Bio-Rad). Reduction in kinase activity upon drug treatment of each sample was confirmed by a CK2 activity assay (CycLex CK2 Kinase Assay Kit (MBL Life Science)) on samples diluted 1:10 in XB. Metaphase arrest of each extract sample after 1h spindle assembly reaction was confirmed by spindle spin down, fixation and imaging as described above.

Proteins from egg and embryo extracts were precipitated with ice-cold acetone and resuspended in ice-cold lysis buffer buffer (8 M urea, 25 mM Tris-HCl pH 8.5, 150 mM NaCl, phosphatase inhibitors (2.5 mM β-glycerophosphate, 1 mM sodium fluoride, 1 mM sodium orthovanadate, 1 mM sodium molybdate) and protease inhibitors (1 mini-Complete EDTA-free tablet per 10 ml lysis buffer; Roche Life Sciences)) and sonicated three times for 15 sec each with intermittent cooling on ice. Lysates were centrifuged at 15,000 x g for 30 minutes at 4°C. A portion was removed to determine protein concentration by BCA assay (Pierce/Thermo Fisher Scientific). The remaining supernatants were reduced with 5 mM DTT at 55°C for 30 minutes, cooled to room temperature, and alkylated with 15 mM iodoacetamide at room temperature, in the dark, for 45 minutes. Alkylation reactions were quenched with an additional 5 mM of DTT for 10 minutes at room temperature then diluted 6-fold with 25 mM Tris pH 8.1 before digestion with trypsin at concentration 1:100 (w/w) (Sigma) at 37°C, overnight. The next day, digests were quenched by acidification with 0.25% TFA (v/v), centrifuged to remove precipitation and desalted with Oasis HLB 60 mg plate (Waters). For proteomics analysis, 40 microgram of peptide digests from each sample were removed. The remaining peptides were lyophilized and stored at -80°C before further analysis.

### Proteomics, phosphoproteomics and data normalization

For proteomics analysis, peptides were labeled with TMTpro reagents (Thermo Fisher Scientific) in 100 mM EPPS pH 8.5/20% ACN at room temperature for 1 hour before a small portion was removed from each sample to determine labeling efficiency. Labeling was confirmed to be at least 95% efficient before quenching with the addition of hydroxylamine to a final concentration of 0.25% for 10 minutes, mixed, acidified with TFA to a pH of about 2, and desalted over an Oasis HLB 10 mg plate (Waters). The desalted multiplex was dried by vacuum centrifugation and separated by offline Pentafluorophenyl (PFP)-based reversed-phase HPLC fractionation as previously described (Grassetti et al., 2017).

For phosphoproteomics analysis, lyophilized digested peptides were subjected to phosphopeptide enrichment with High Select Fe-NTA Phosphopeptide Enrichment Kit (Thermo Fisher Scientific) following the manufacturer’s instructions and desalted over an Oasis HLB 10 mg plate (Waters). Phosphopeptides were then labeled with TMTpro reagents and offline separated as described above.

TMT-labeled peptides were analyzed on an Orbitrap Lumos mass spectrometer (ThermoScientific) equipped with an Easy-nLC 1200 (ThermoScientific). The raw data files were searched using COMET (Eng et al., 2013) with a static mass of 304.2071 Da on peptide N-termini and lysines, 57.02146 Da on cysteines, and a variable mass of 15.99491 Da on methionines and for phosphorylation 79.96633 Da on serines, threonines, and tyrosines against the target-decoy version (Elias and Gygi, 2007) of the annotated *Xenopus laevis* proteome database (Xenbase, v9.2) (Fisher et al., 2023) and filtered to a <1% FDR at the peptide level. Quantification of LC-MS/MS spectra was performed using in-house developed software. The probability of phosphorylation site localization was determined by PhosphoRS (Taus et al., 2011). For proteomics analysis, peptide intensities were adjusted based on total TMT reporter ion intensity in each channel across all samples, and log2 transformed. For phosphoproteomics analysis, peptide intensities were adjusted based on total TMT reporter ion intensity in each channel across all samples (or for phase normalized data across meiotic samples or mitotic samples, respectively), and log2 transformed. P-values were calculated using a two-tailed Student’s t-test in Perseus (Tyanova et al., 2016).

### GO enrichment and Network Analysis

For analysis, proteomic and phosphoproteomic data from the two experimental repeats (total peptide count >1 filtered) was pooled. Data was annotated with human orthologs using the mapping to the *X. laevis* v9.2 genome in the files available at https://wuehr.scholar.princeton.edu/protein-concentrations-xenopus-egg (Wühr et al., 2014). GO enrichment analysis was carried out on human orthologs using PANTHER version 19.0 (Thomas et al., 2022) with default parameters. Network analysis was carried out on human orthologs using STRING version 12.0 (Szklarczyk et al., 2019) using full STRING network annotations with high confidence (interaction score ≥ 0.7). Edge color indicates data source with known interactions shown in cyan (from curated databases) and magenta (experimentally determined), predicted interactions shown in emerald green (gene neighbourhood), red (gene fusions) and royal blue (gene co-occurrence), and lime green (textmining), black (co-expression) and lilac (protein homology). K-means clustering with default parameters was used to define functional clusters.

### Statistical tests

Mann-Whitney and t-tests (as indicated in figure legends) were carried out using Prism 9.0.

## Supplemental material

There are 6 supplementary figures and 8 supplementary tables:

Figure S1 – Spindle length changes dramatically at the meiosis to mitosis transition in *Ciona* and *Xenopus* though cell size does not.

Figure S2 – Effects of small molecules in screen on meiotic spindle morphology in *Ciona* oocytes and effects of CK2 inhibition on ectopic aster formation in *X. laevis* extracts.

Figure S3 – CK2 activity is reduced by CK2 inhibition in *X. laevis* extract proteomics and phosphoproteomics samples.

Figure S4 - Proteomic changes between meiosis and mitosis and upon CK2 inhibition.

Figure S5 - Phosphoproteomic changes between meiosis and mitosis and upon CK2 inhibition.

Figure S6 - Inhibition of RanGTP-mediated spindle assembly pathway reduces TPX2 and tubulin intensity at spindle poles in *X. laevis* egg extracts.

Table S1 – Summary of proteomic data

Table S2 – Summary of phosphoproteomic data

Table S3 – GO enrichment analysis results – proteins enriched in meiosis relative to mitosis

Table S4 – Microtubule binding proteins enriched in meiosis relative to mitosis

Table S5 – GO enrichment analysis results – proteins enriched in mitosis relative to meiosis

Table S6 – GO enrichment analysis results – proteins with phosphosites enriched in mitosis

Table S7 – Proteins in Spindle GO category (GO:0005819) with phosphosites enriched in mitosis relative to meiosis

Table S8 – Candidate proteins annotated with localization data

## Acknowledgements

We thank Matthew Swaffer, Coral Zhou, Xiao Liu, Gabriel Cavin-Meza and Cordell Clark for feedback on the manuscript, Yasaswini Sampathkumar for importin-β(71-876) protein, Michael Levine and Laurence Lemaire for training and advice on *Ciona* experiments and Marina Crowder for advice. This work was supported by NIH MIRA grant R35GM118183 and the Flora Lamson Hewlett Chair in Biochemistry to RH, an EMBO long term postdoctoral fellowship to HC and NIGMS grant R35GM119455 to AK.

---

**Figure S1.**
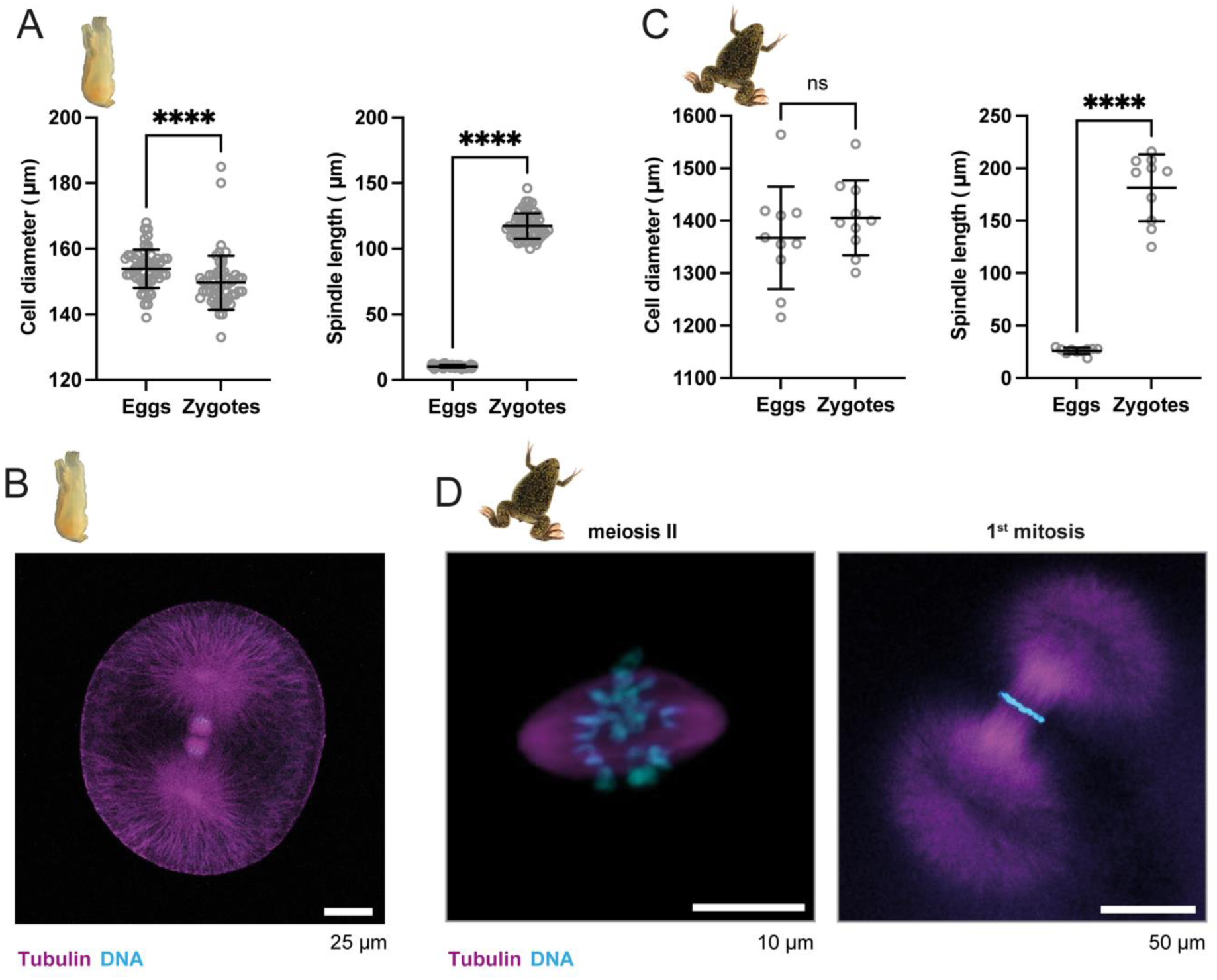
Spindle length changes dramatically at the meiosis to mitosis transition in *Ciona* and *Xenopus* though cell size does not. **(A)** *C. robusta* cell diameter and metaphase spindle length (eggs n=58, zygotes n=58) measurements. Statistical significance determined by two-tailed Mann Whitney test (**** = p < 0.00005). Line indicates mean and error bars indicate one standard deviation above and below the mean. **(B)** Image of immunofluorescence of a representative *C. robusta* zygote in anaphase of the first mitosis. Tubulin (magenta), DNA (cyan). Single z-slice through center of spindle shown. **(C)** *X. laevis* cell diameter and metaphase spindle length (eggs n=10, zygotes n=10) measurements. Statistical significance determined by two-tailed Mann Whitney test (ns = not significant, **** = p < 0.00005). Line indicates mean and error bars indicate one standard deviation above and below the mean. **(D)** Higher magnification images of immunofluorescence of spindles from representative *X. laevis* egg in metaphase of meiosis II (left panel) and zygote in metaphase of the first mitosis (right panel) shown in Figure 1D. Tubulin (magenta), DNA (cyan). Single z-slice through center of spindle shown.

**Figure S2.**
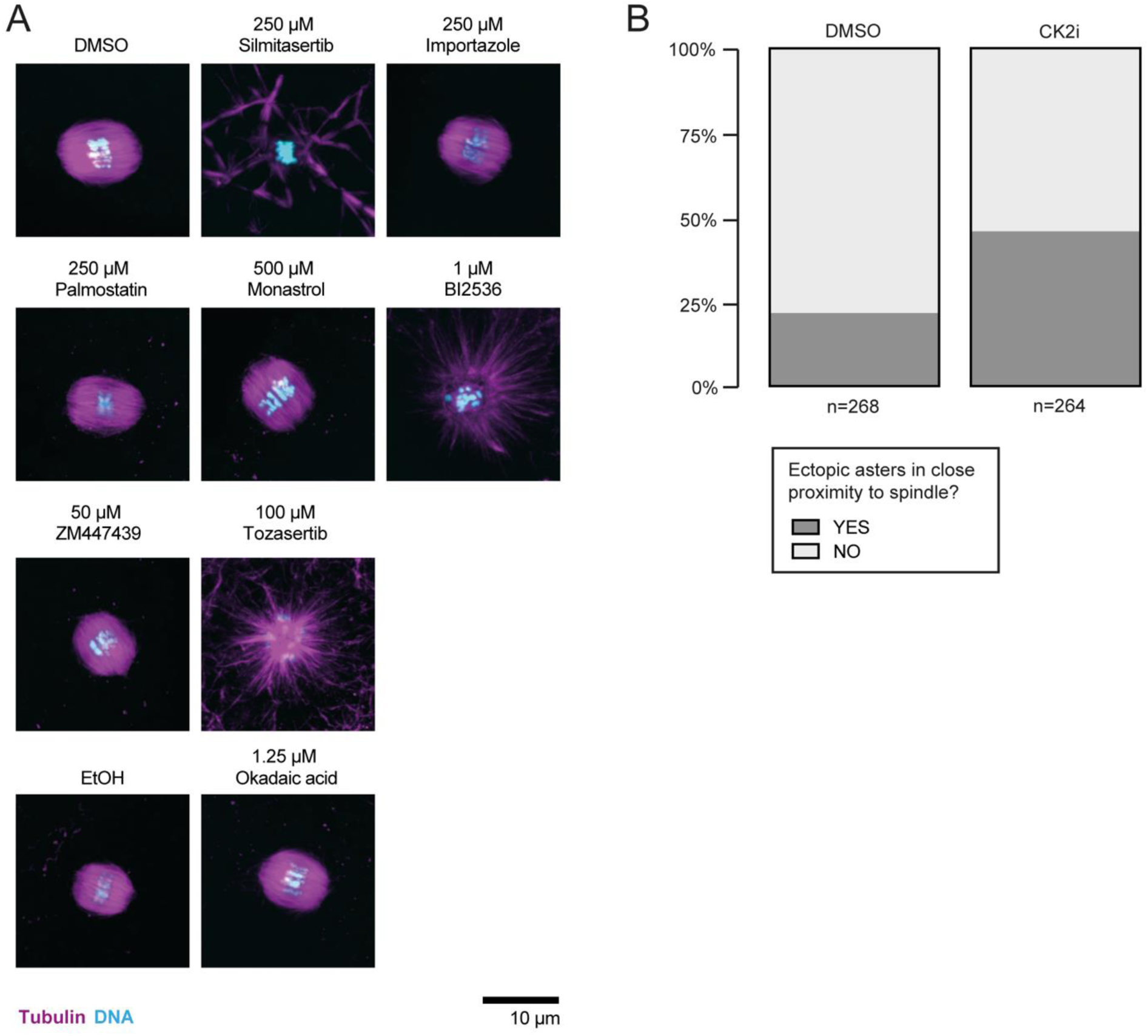
Effects of small molecules in screen on meiotic spindle morphology in *Ciona* oocytes and effects of CK2 inhibition on ectopic aster formation in *X. laevis* extracts. **(A)** Images of immunofluorescence of representative *C. robusta* oocytes treated for 1 hour with DMSO, 250 µM silmitasertib (CK2i), 250 µM RanGTP-importin β interaction inhibitor importazole, 250 µM APT1 inhibitor palmostatin, 500 µM Eg5 inhibitor monastrol, 1 µM Plk1 inhibitor BI2536, 50 µM Aurora A and B inhibitor ZM447439, 100 µM pan-aurora inhibitor tozasertib, EtOH or 1.25 µM PP1 and PP2A inhibitor okadaic acid as indicated. Okadaic acid was in EtOH and all other drugs were in DMSO. Maximum intensity projections shown. Tubulin (magenta), DNA (cyan). **(B)** Percentage of bipolar spindles displaying ectopic microtubule asters in proximity to the spindle in metaphasic CSF *X. laevis* egg extract in the presence of 50 µM CK2i or DMSO as indicated. Data from four independent extract experiments pooled (n = 264 spindles (CK2i), n = 268 spindles (DMSO)); n ≥ 40 spindles per condition for each extract.

**Figure S3.**
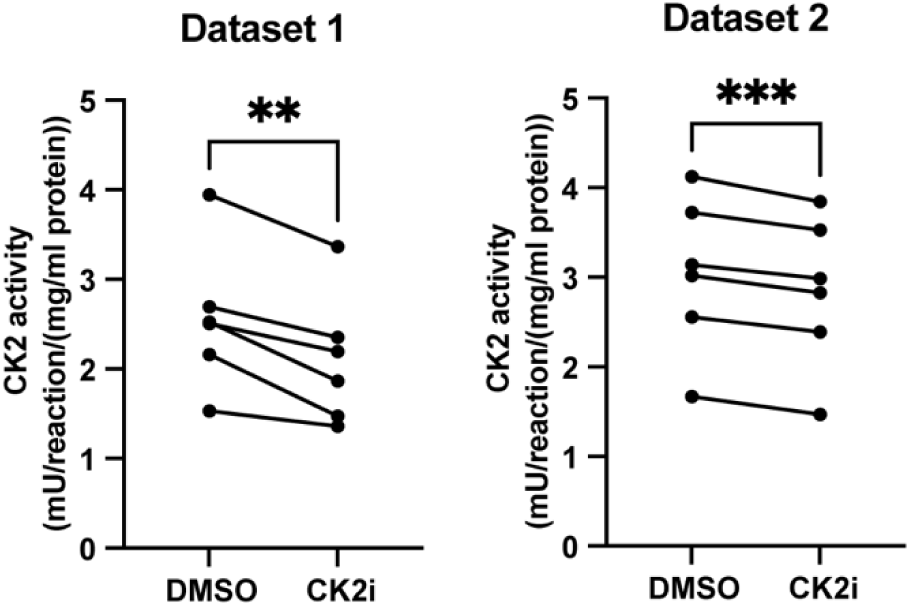
CK2 activity is reduced by CK2 inhibition in *X. laevis* extract proteomics and phosphoproteomics samples. CK2 activity of 6 extracts (3 meiotic and 3 mitotic) used in proteomics and phosphoproteomics experiments in control and CK2i treated samples for each dataset. Statistical significance determined by paired t test (** = p < 0.005, *** = p < 0.0005).

**Figure S4.**
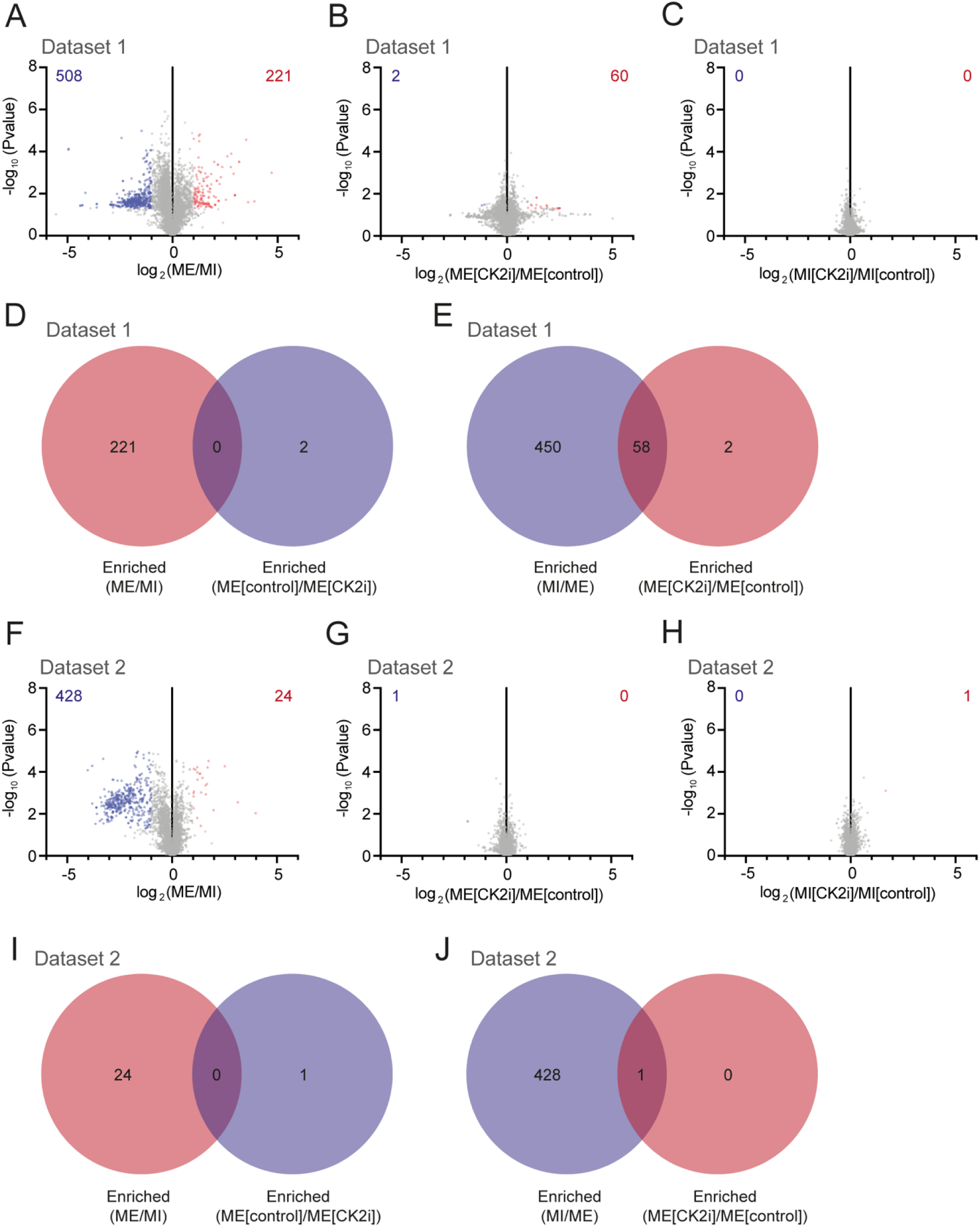
Proteomic changes between meiosis and mitosis and upon CK2 inhibition. **(A-C)** Volcano plots of dataset 1 log_2_ ratio (protein abundance) versus -log_10_ p value for individual proteins in Meiosis compared to Mitosis (ME/MI) **(A)**, Meiosis in the presence of CK2i compared to Meiosis in the absence of CK2i **(B)** and Mitosis in the presence of CK2i compared to Mitosis in the absence of CK2i **(C)** respectively. Red points indicate significantly enriched (log_2_ ratio >1 and p value <0.05) proteins (total number indicated in red). Blue points indicate significantly depleted (log_2_ ratio <-1 and p value <0.05) proteins (total number indicated in blue) **(D)** Venn diagram indicating number of unique proteins in dataset 1 in the categories enriched in Meiosis relative to Mitosis and enriched in Meiosis in the absence of CK2i compared to Meiosis in the presence of CK2i and in the overlap between these two categories **(E)** Venn diagram indicating number of unique proteins in dataset 1 in the categories enriched in Mitosis relative to Meiosis and enriched in Meiosis in the presence of CK2i compared to Meiosis in the absence of CK2i and in the overlap between these two categories **(F-H)** Volcano plots of dataset 2 log_2_ ratio (protein abundance) versus -log_10_ p value for individual proteins in Meiosis compared to Mitosis (ME/MI) **(F)**, Meiosis in the presence of CK2i compared to Meiosis in the absence of CK2i **(G)** and Mitosis in the presence of CK2i compared to Mitosis in the absence of CK2i **(H)** respectively. Red points indicate significantly enriched (log_2_ ratio >1 and p value <0.05) proteins (total number indicated in red). Blue points indicate significantly depleted (log_2_ ratio <-1 and p value <0.05) proteins (total number indicated in blue) **(I)** Venn diagram indicating number of unique proteins in dataset 2 in the categories enriched in Meiosis relative to Mitosis and enriched in Meiosis in the absence of CK2i compared to Meiosis in the presence of CK2i and in the overlap between these two categories **(J)** Venn diagram indicating number of unique proteins in dataset 2 in the categories enriched in Mitosis relative to Meiosis and enriched in Meiosis in the presence of CK2i compared to Meiosis in the absence of CK2i and in the overlap between these two categories.

**Figure S5.**
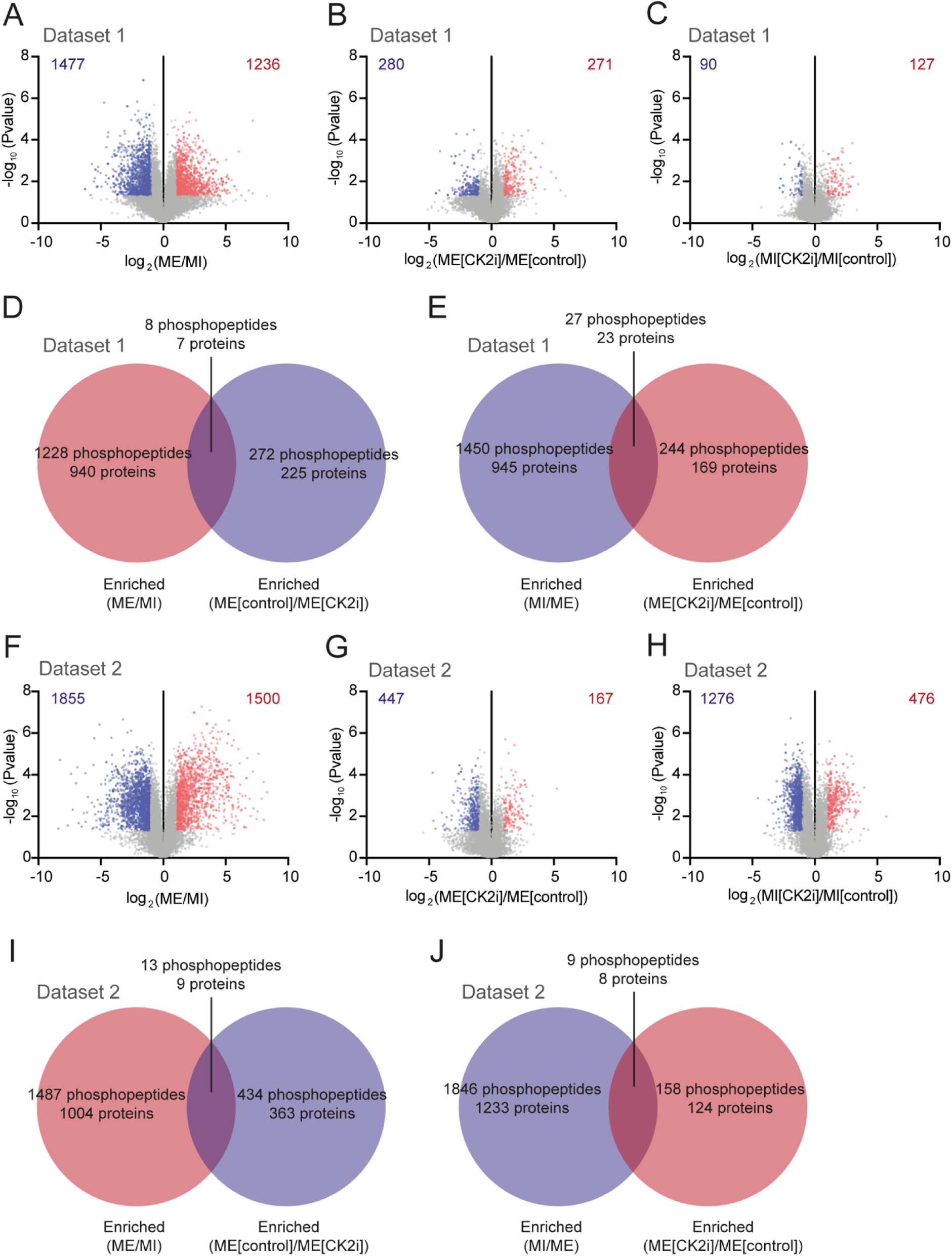
Phosphoproteomic changes between meiosis and mitosis and upon CK2 inhibition. **(A-C)** Volcano plots of dataset 1 log_2_ ratio (phosphopeptide abundance) versus -log_10_ p value for individual phosphopeptides in Meiosis compared to Mitosis (ME/MI) **(A)**, Meiosis in the presence of CK2i compared to Meiosis in the absence of CK2i **(B)** and Mitosis in the presence of CK2i compared to Mitosis in the absence of CK2i **(C)** respectively. Red points indicate significantly enriched (log_2_ ratio >1 and p value <0.05) phosphopeptides (total number indicated in red). Blue points indicate significantly depleted (log_2_ ratio <-1 and p value <0.05) phosphopeptides (total number indicated in blue) **(D)** Venn diagram indicating number of unique phosphopeptides and proteins containing phosphopeptides in dataset 1 in the categories enriched in Meiosis relative to Mitosis and enriched in Meiosis in the absence of CK2i compared to Meiosis in the presence of CK2i and in the overlap between these two categories that do not exhibit protein level changes in the same directions **(E)** Venn diagram indicating number of unique phosphopepetides and proteins containing phosphopeptides in dataset 1 in the categories enriched in Mitosis relative to Meiosis and enriched in Meiosis in the presence of CK2i compared to Meiosis in the absence of CK2i and in the overlap between these two categories that do not exhibit protein level changes in the same directions **(F-H)** Volcano plots of dataset 2 log_2_ ratio (phosphopeptide abundance) versus -log_10_ p value for individual phosphopeptides in Meiosis compared to Mitosis (ME/MI) **(F)**, Meiosis in the presence of CK2i compared to Meiosis in the absence of CK2i **(G)** and Mitosis in the presence of CK2i compared to Mitosis in the absence of CK2i **(H)** respectively. Red points indicate significantly enriched (log_2_ ratio >1 and p value <0.05) phosphopeptides (total number indicated in red). Blue points indicate significantly depleted (log_2_ ratio <-1 and p value <0.05) phosphopeptides (total number indicated in blue) **(I)** Venn diagram indicating number of unique phosphopeptides and proteins containing phosphopeptides in dataset 2 in the categories enriched in Meiosis relative to Mitosis and enriched in Meiosis in the absence of CK2i compared to Meiosis in the presence of CK2i and in the overlap between these two categories that do not exhibit protein level changes in the same directions **(J)** Venn diagram indicating number of unique phosphopepetides and proteins containing phosphopeptides in dataset 2 in the categories enriched in Mitosis relative to Meiosis and enriched in Meiosis in the presence of CK2i compared to Meiosis in the absence of CK2i and in the overlap between these two categories that do not exhibit protein level changes in the same directions.

**Figure S6.**
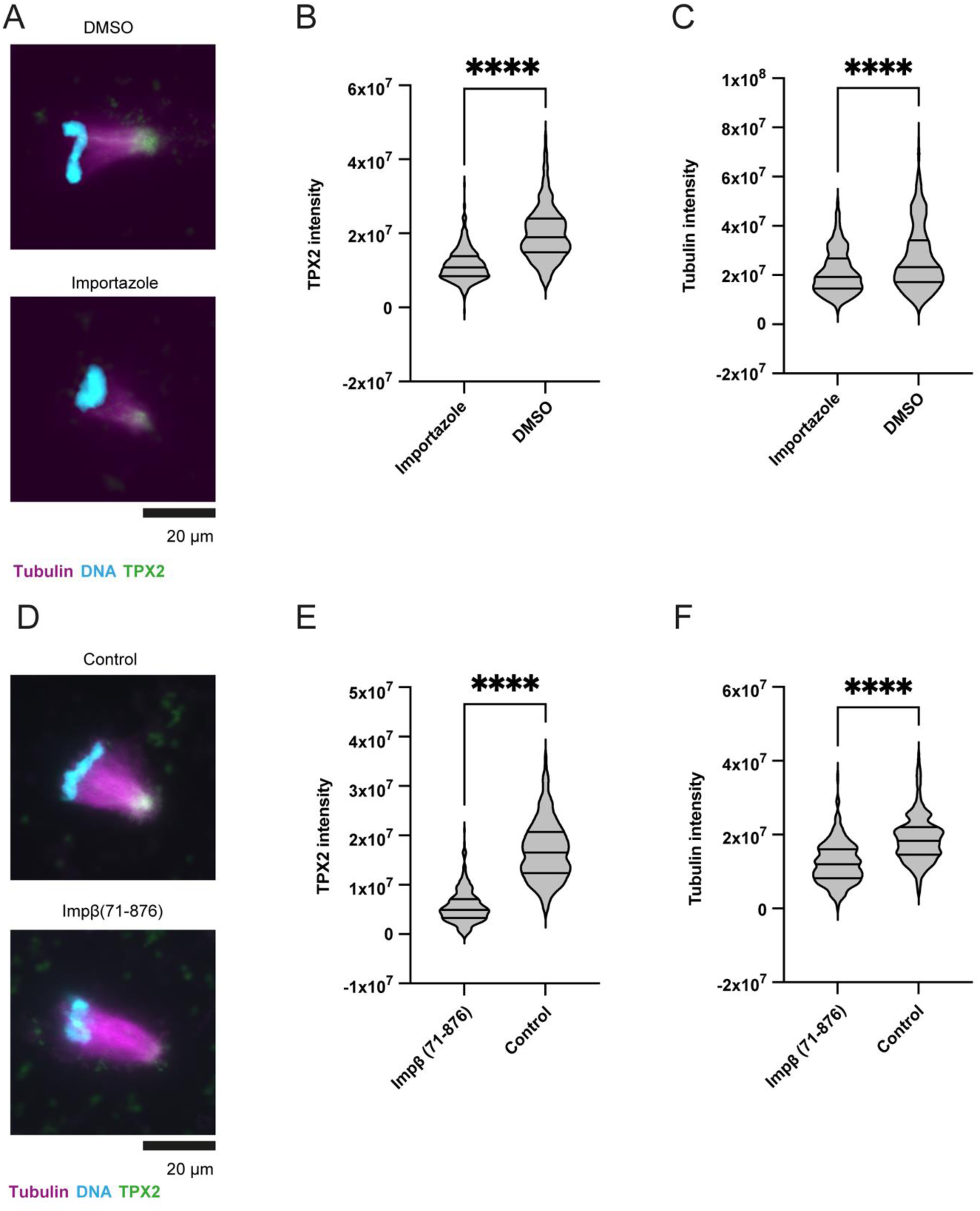
Inhibition of RanGTP-mediated spindle assembly pathway reduces TPX2 and tubulin intensity at spindle poles in *X. laevis* egg extracts. **(A)** Representative images of half spindles from spindle assembly reactions carried out in the presence of DMSO or 300 µM importazole as indicated. β tubulin (magenta), DNA (cyan), TPX2 (green). **(B-C)** Violin plots of TPX2 intensity (B) or β tubulin intensity (C) of half spindles from experiment described in (A). n≥420 half spindles per condition (from three cytoplasmic extracts (n≥133 per extract for each condition)). **(D)** Representative images of half spindles from spindle assembly reactions carried out in the presence of XB (control) or 2.5 µM Importin-β(71-876) as indicated. β tubulin (magenta), DNA (cyan), TPX2 (green). **(E-F)** Violin plot of TPX2 intensity (E) or β tubulin intensity (F) of half spindles from experiment described in (B). n≥329 half spindles per condition (from three cytoplasmic extracts (n≥92 per extract for each condition)). Statistical significance determined by two-tailed Mann Whitney test (**** = p < 0.00005). Lines indicate median and upper and lower quartiles. Data is a subset of data plotted in Figure 6.

## Notes

### Competing Interest Statement

The authors have declared no competing interest.

