## Supplementary Table S3 for "Spindle morphology changes between meiosis and mitosis driven by CK2 regulation of the Ran pathway"

| GO category | | Enriched in meiosis | | | | |
|---|---|---|---|---|---|---|
| | Background | Number of proteins | Expected number proteins | Fold enrichment | Raw p value | False discovery rate |
| ***PANTHER GO-Slim Molecular Function*** | | | | | | |
| structural constituent of cytoskeleton (GO:0005200) | 21 | 5 | 0.54 | 9.23 | 1.53E-04 | 1.37E-02 |
| ubiquitin protein ligase activity (GO:0061630) | 50 | 10 | 1.29 | 7.76 | 3.76E-07 | 2.03E-04 |
| ubiquitin-like protein ligase activity (GO:0061659) | 55 | 10 | 1.42 | 7.05 | 9.60E-07 | 2.59E-04 |
| microtubule binding (GO:0008017) | 42 | 6 | 1.08 | 5.54 | 6.33E-04 | 3.79E-02 |
| ubiquitin-protein transferase activity (GO:0004842) | 79 | 11 | 2.04 | 5.4 | 4.25E-06 | 7.63E-04 |
| ubiquitin-like protein transferase activity (GO:0019787) | 87 | 11 | 2.24 | 4.9 | 1.11E-05 | 1.49E-03 |
| aminoacyltransferase activity (GO:0016755) | 88 | 11 | 2.27 | 4.85 | 1.24E-05 | 1.33E-03 |
| acyltransferase activity (GO:0016746) | 123 | 11 | 3.17 | 3.47 | 2.79E-04 | 2.15E-02 |
| transferase activity (GO:0016740) | 439 | 23 | 11.32 | 2.03 | 7.13E-04 | 3.84E-02 |
| ***PANTHER GO-Slim Cellular Component*** | | | | | | |
| intermediate filament cytoskeleton (GO:0045111) | 10 | 4 | 0.26 | 15.51 | 7.81E-05 | 1.78E-02 |
| intermediate filament (GO:0005882) | 10 | 4 | 0.26 | 15.51 | 7.81E-05 | 1.19E-02 |
| extracellular space (GO:0005615) | 72 | 8 | 1.86 | 4.31 | 4.61E-04 | 3.50E-02 |
| extracellular region (GO:0005576) | 76 | 8 | 1.96 | 4.08 | 6.67E-04 | 4.34E-02 |
| ***PANTHER GO-Slim Biological Process*** | | | | | | |
| production of molecular mediator of immune response (GO:0002440) | 6 | 3 | 0.15 | 19.39 | 3.16E-04 | 4.94E-02 |
| intermediate filament organization (GO:0045109) | 13 | 4 | 0.34 | 11.93 | 2.50E-04 | 4.28E-02 |
| innate immune response (GO:0045087) | 14 | 4 | 0.36 | 11.08 | 3.44E-04 | 4.96E-02 |
| defense response to symbiont (GO:0140546) | 18 | 5 | 0.46 | 10.77 | 6.84E-05 | 4.28E-02 |
| immune response (GO:0006955) | 29 | 8 | 0.75 | 10.7 | 4.16E-07 | 7.82E-04 |
| defense response to other organism (GO:0098542) | 19 | 5 | 0.49 | 10.2 | 9.09E-05 | 4.27E-02 |
| response to external biotic stimulus (GO:0043207) | 22 | 5 | 0.57 | 8.81 | 1.93E-04 | 7.27E-02 |
| response to other organism (GO:0051707) | 22 | 5 | 0.57 | 8.81 | 1.93E-04 | 6.05E-02 |
| response to biotic stimulus (GO:0009607) | 22 | 5 | 0.57 | 8.81 | 1.93E-04 | 5.19E-02 |
| biological process involved in interspecies interaction between organisms (GO:0044419) | 22 | 5 | 0.57 | 8.81 | 1.93E-04 | 4.54E-02 |
| defense response (GO:0006952) | 23 | 5 | 0.59 | 8.43 | 2.42E-04 | 4.55E-02 |
| immune system process (GO:0002376) | 41 | 8 | 1.06 | 7.57 | 7.16E-06 | 6.72E-03 |
| protein ubiquitination (GO:0016567) | 81 | 9 | 2.09 | 4.31 | 2.02E-04 | 4.21E-02 |

### Table S3 – GO enrichment analysis results – proteins enriched in meiosis relative to mitosis
