## Supplementary Table S4 for "Spindle morphology changes between meiosis and mitosis driven by CK2 regulation of the Ran pathway"

| Gene ID | UNIPROT ID | Gene Name | Gene symbol |
|---|---|---|---|
| HUMAN\|HGNC=17429\|UniProtKB=Q9P2M7 | Q9P2M7 | Cingulin | CGN |
| HUMAN\|HGNC=16691\|UniProtKB=Q9UGJ1 | Q9UGJ1 | Gamma-tubulin complex component 4 | TUBGCP4 |
| HUMAN\|HGNC=6391\|UniProtKB=Q14807 | Q14807 | Kinesin-like protein KIF22 | KIF22 |
| HUMAN\|HGNC=6892\|UniProtKB=Q9UPY8 | Q9UPY8 | Microtubule-associated protein RP_EB family member 3 | MAPRE3 |
| HUMAN\|HGNC=29789\|UniProtKB=Q9ULD2 | Q9ULD2 | Microtubule-associated tumor suppressor 1 | MTUS1 |
| HUMAN\|HGNC=9341\|UniProtKB=O43663 | O43663 | Protein regulator of cytokinesis 1 | PRC1 |

**Table S4 – Microtubule binding proteins enriched in meiosis relative to mitosis**
