## Supplementary Table S5 for "Spindle morphology changes between meiosis and mitosis driven by CK2 regulation of the Ran pathway"

| GO category | Background | Enriched in mitosis | | | | |
| --- | --- | --- | --- | --- | --- | --- |
| | | Number of proteins | Expected number proteins | Fold enrichment | Raw p value | False discovery rate |
| ***PANTHER GO-Slim Molecular Function*** | | | | | | |
| palmitoyltransferase activity (GO:0016409) | 3 | 3 | 0.24 | 12.34 | 5.28E-04 | 1.67E-02 |
| UDP-galactosyltransferase activity (GO:0035250) | 3 | 3 | 0.24 | 12.34 | 5.28E-04 | 1.58E-02 |
| protein disulfide isomerase activity (GO:0003756) | 8 | 7 | 0.65 | 10.8 | 1.62E-07 | 2.18E-05 |
| galactosyltransferase activity (GO:0008378) | 5 | 4 | 0.41 | 9.87 | 1.99E-04 | 8.24E-03 |
| O-acyltransferase activity (GO:0008374) | 5 | 4 | 0.41 | 9.87 | 1.99E-04 | 7.65E-03 |
| glucosyltransferase activity (GO:0046527) | 8 | 6 | 0.65 | 9.25 | 6.62E-06 | 7.14E-04 |
| cholesterol binding (GO:0015485) | 4 | 3 | 0.32 | 9.25 | 1.99E-03 | 3.96E-02 |
| hexosyltransferase activity (GO:0016758) | 26 | 15 | 2.11 | 7.12 | 1.09E-10 | 5.86E-08 |
| UDP-glycosyltransferase activity (GO:0008194) | 14 | 7 | 1.13 | 6.17 | 4.51E-05 | 2.70E-03 |
| oxidoreductase activity, acting on paired donors, with incorporation or reduction of molecular oxygen (GO:0016705) | 20 | 9 | 1.62 | 5.55 | 1.03E-05 | 9.21E-04 |
| heme binding (GO:0020037) | 14 | 6 | 1.13 | 5.29 | 4.66E-04 | 1.67E-02 |
| monooxygenase activity (GO:0004497) | 12 | 5 | 0.97 | 5.14 | 1.67E-03 | 3.75E-02 |
| glycosyltransferase activity (GO:0016757) | 41 | 17 | 3.32 | 5.12 | 4.80E-09 | 1.29E-06 |
| tetrapyrrole binding (GO:0046906) | 15 | 6 | 1.22 | 4.94 | 7.24E-04 | 2.05E-02 |
| oxidoreductase activity, acting on a sulfur group of donors (GO:0016667) | 22 | 7 | 1.78 | 3.93 | 1.27E-03 | 3.10E-02 |
| calcium ion binding (GO:0005509) | 34 | 9 | 2.76 | 3.27 | 1.15E-03 | 2.96E-02 |
| acyltransferase activity, transferring groups other than amino-acyl groups (GO:0016747) | 31 | 8 | 2.51 | 3.18 | 2.58E-03 | 4.96E-02 |
| isomerase activity (GO:0016853) | 59 | 14 | 4.78 | 2.93 | 1.86E-04 | 8.35E-03 |
| transmembrane transporter activity (GO:0022857) | 85 | 16 | 6.89 | 2.32 | 1.78E-03 | 3.85E-02 |
| oxidoreductase activity (GO:0016491) | 176 | 32 | 14.26 | 2.24 | 1.25E-05 | 9.63E-04 |
| transporter activity (GO:0005215) | 95 | 17 | 7.7 | 2.21 | 1.67E-03 | 3.91E-02 |
| ***PANTHER GO-Slim Cellular Component*** | | | | | | |
| EMC complex (GO:0072546) | 7 | 7 | 0.57 | 12.34 | 2.17E-08 | 7.63E-07 |
| rough endoplasmic reticulum membrane (GO:0030867) | 3 | 3 | 0.24 | 12.34 | 5.28E-04 | 7.30E-03 |
| oligosaccharyltransferase complex (GO:0008250) | 5 | 5 | 0.41 | 12.34 | 3.41E-06 | 8.63E-05 |
| rough endoplasmic reticulum (GO:0005791) | 3 | 3 | 0.24 | 12.34 | 5.28E-04 | 7.08E-03 |
| endoplasmic reticulum protein-containing complex (GO:0140534) | 31 | 26 | 2.51 | 10.35 | 2.07E-24 | 1.35E-22 |
| endoplasmic reticulum lumen (GO:0005788) | 5 | 4 | 0.41 | 9.87 | 1.99E-04 | 3.36E-03 |
| endoplasmic reticulum-Golgi intermediate compartment (GO:0005793) | 17 | 13 | 1.38 | 9.43 | 9.33E-12 | 4.25E-10 |
| endoplasmic reticulum subcompartment (GO:0098827) | 73 | 53 | 5.92 | 8.96 | 3.37E-43 | 5.12E-41 |
| nuclear outer membrane-endoplasmic reticulum membrane network (GO:0042175) | 72 | 52 | 5.84 | 8.91 | 3.48E-42 | 3.97E-40 |
| endoplasmic reticulum membrane (GO:0005789) | 72 | 52 | 5.84 | 8.91 | 3.48E-42 | 3.18E-40 |
| endoplasmic reticulum (GO:0005783) | 164 | 115 | 13.29 | 8.65 | 1.17E-93 | 5.33E-91 |
| COPII-coated ER to Golgi transport vesicle (GO:0030134) | 20 | 13 | 1.62 | 8.02 | 2.42E-10 | 1.00E-08 |
| organelle subcompartment (GO:0031984) | 95 | 57 | 7.7 | 7.4 | 2.53E-39 | 1.92E-37 |
| Golgi membrane (GO:0000139) | 14 | 6 | 1.13 | 5.29 | 4.66E-04 | 6.85E-03 |
| endomembrane system (GO:0012505) | 428 | 142 | 34.69 | 4.09 | 6.02E-59 | 1.37E-56 |
| coated vesicle (GO:0030135) | 45 | 13 | 3.65 | 3.56 | 3.49E-05 | 7.23E-04 |
| organelle membrane (GO:0031090) | 294 | 62 | 23.83 | 2.6 | 2.81E-13 | 1.43E-11 |
| Golgi apparatus (GO:0005794) | 99 | 20 | 8.02 | 2.49 | 9.59E-05 | 1.75E-03 |
| membrane (GO:0016020) | 661 | 110 | 53.57 | 2.05 | 1.86E-15 | 1.06E-13 |
| membrane protein complex (GO:0098796) | 198 | 30 | 16.05 | 1.87 | 7.06E-04 | 9.20E-03 |
| intracellular membrane-bounded organelle (GO:0043231) | 1783 | 189 | 144.51 | 1.31 | 7.78E-07 | 2.37E-05 |
| membrane-bounded organelle (GO:0043227) | 1825 | 192 | 147.92 | 1.3 | 8.70E-07 | 2.48E-05 |
| ***PANTHER GO-Slim Biological Process*** | | | | | | |
| endoplasmic reticulum tubular network organization (GO:0071786) | 5 | 5 | 0.41 | 12.34 | 3.41E-06 | 4.00E-04 |
| protein N-linked glycosylation via asparagine (GO:0018279) | 5 | 5 | 0.41 | 12.34 | 3.41E-06 | 3.76E-04 |
| protein N-linked glycosylation (GO:0006487) | 12 | 8 | 0.97 | 8.23 | 6.39E-07 | 1.00E-04 |
| glycolipid biosynthetic process (GO:0009247) | 6 | 4 | 0.49 | 8.23 | 5.58E-04 | 2.62E-02 |
| endoplasmic reticulum organization (GO:0007029) | 15 | 10 | 1.22 | 8.23 | 2.25E-08 | 7.04E-06 |
| membrane lipid biosynthetic process (GO:0046467) | 9 | 6 | 0.73 | 8.23 | 1.85E-05 | 1.24E-03 |
| membrane lipid metabolic process (GO:0006643) | 10 | 6 | 0.81 | 7.4 | 4.31E-05 | 2.79E-03 |
| liposaccharide metabolic process (GO:1903509) | 7 | 4 | 0.57 | 7.05 | 1.22E-03 | 4.77E-02 |
| glycolipid metabolic process (GO:0006664) | 7 | 4 | 0.57 | 7.05 | 1.22E-03 | 4.67E-02 |
| SRP-dependent cotranslational protein targeting to membrane (GO:0006614) | 7 | 4 | 0.57 | 7.05 | 1.22E-03 | 4.58E-02 |
| cotranslational protein targeting to membrane (GO:0006613) | 7 | 4 | 0.57 | 7.05 | 1.22E-03 | 4.49E-02 |
| protein localization to endoplasmic reticulum (GO:0070972) | 15 | 8 | 1.22 | 6.58 | 6.68E-06 | 5.97E-04 |
| protein glycosylation (GO:0006486) | 18 | 9 | 1.46 | 6.17 | 3.44E-06 | 3.59E-04 |
| macromolecule glycosylation (GO:0043413) | 18 | 9 | 1.46 | 6.17 | 3.44E-06 | 3.40E-04 |
| glycosylation (GO:0070085) | 20 | 10 | 1.62 | 6.17 | 9.55E-07 | 1.38E-04 |
| endoplasmic reticulum to Golgi vesicle-mediated transport (GO:0006888) | 36 | 17 | 2.92 | 5.83 | 3.98E-10 | 3.74E-07 |
| response to endoplasmic reticulum stress (GO:0034976) | 30 | 14 | 2.43 | 5.76 | 1.80E-08 | 6.76E-06 |
| protein targeting to ER (GO:0045047) | 13 | 6 | 1.05 | 5.69 | 2.85E-04 | 1.49E-02 |
| establishment of protein localization to endoplasmic reticulum (GO:0072599) | 13 | 6 | 1.05 | 5.69 | 2.85E-04 | 1.45E-02 |
| long-chain fatty acid metabolic process (GO:0001676) | 11 | 5 | 0.89 | 5.61 | 1.04E-03 | 4.26E-02 |
| Golgi organization (GO:0007030) | 22 | 10 | 1.78 | 5.61 | 2.88E-06 | 3.87E-04 |
| glycoprotein biosynthetic process (GO:0009101) | 21 | 9 | 1.7 | 5.29 | 1.67E-05 | 1.20E-03 |
| ERAD pathway (GO:0036503) | 21 | 9 | 1.7 | 5.29 | 1.67E-05 | 1.16E-03 |
| intracellular calcium ion homeostasis (GO:0006874) | 14 | 6 | 1.13 | 5.29 | 4.66E-04 | 2.24E-02 |
| glycoprotein metabolic process (GO:0009100) | 27 | 11 | 2.19 | 5.03 | 3.38E-06 | 4.24E-04 |
| calcium ion homeostasis (GO:0055074) | 15 | 6 | 1.22 | 4.94 | 7.24E-04 | 3.09E-02 |

| | | | | | | |
|---|---|---|---|---|---|---|
| endomembrane system organization (GO:0010256) | 53 | 21 | 4.3 | 4.89 | 1.94E-10 | 3.64E-07 |
| response to organonitrogen compound (GO:0010243) | 35 | 12 | 2.84 | 4.23 | 1.00E-05 | 7.86E-04 |
| response to nitrogen compound (GO:1901698) | 35 | 12 | 2.84 | 4.23 | 1.00E-05 | 7.54E-04 |
| protein folding (GO:0006457) | 65 | 22 | 5.27 | 4.18 | 2.46E-09 | 1.54E-06 |
| Golgi vesicle transport (GO:0048193) | 72 | 21 | 5.84 | 3.6 | 1.11E-07 | 2.61E-05 |
| protein maturation (GO:0051604) | 95 | 27 | 7.7 | 3.51 | 2.99E-09 | 1.40E-06 |
| response to organic substance (GO:0010033) | 54 | 13 | 4.38 | 2.97 | 2.70E-04 | 1.45E-02 |
| response to chemical (GO:0042221) | 92 | 19 | 7.46 | 2.55 | 1.05E-04 | 6.57E-03 |
| lipid metabolic process (GO:0006629) | 117 | 22 | 9.48 | 2.32 | 1.77E-04 | 1.04E-02 |

### Table S5 – GO enrichment analysis results – proteins enriched in mitosis relative to meiosis
