## Supplementary Table S6 for "Spindle morphology changes between meiosis and mitosis driven by CK2 regulation of the Ran pathway"

| GO category | Background | Enriched in mitosis | | | | |
| --- | --- | --- | --- | --- | --- | --- |
| | | Number of proteins | Expected number proteins | Fold enrichment | Raw p value | False discovery rate |
| ***PANTHER GO-Complete Molecular Function*** | | | | | | |
| DNA binding (GO:0003677) | 483 | 185 | 144.96 | 1.28 | 1.69E-05 | 3.33E-02 |
| ***PANTHER GO-Complete Cellular Component*** | | | | | | |
| mitotic spindle (GO:0072686) | 69 | 34 | 20.71 | 1.64 | 7.48E-04 | 4.91E-02 |
| nuclear outer membrane-endoplasmic reticulum membrane network (GO:0042175) | 141 | 68 | 42.32 | 1.61 | 2.94E-06 | 5.52E-04 |
| endoplasmic reticulum subcompartment (GO:0098827) | 139 | 67 | 41.72 | 1.61 | 3.86E-06 | 5.64E-04 |
| endoplasmic reticulum membrane (GO:0005789) | 139 | 67 | 41.72 | 1.61 | 3.86E-06 | 5.07E-04 |
| spindle (GO:0005819) | 148 | 64 | 44.42 | 1.44 | 4.31E-04 | 2.98E-02 |
| organelle subcompartment (GO:0031984) | 194 | 81 | 58.22 | 1.39 | 3.20E-04 | 2.63E-02 |
| endoplasmic reticulum (GO:0005783) | 216 | 89 | 64.82 | 1.37 | 2.57E-04 | 2.25E-02 |
| nuclear body (GO:0016604) | 264 | 105 | 79.23 | 1.33 | 3.80E-04 | 2.77E-02 |
| nucleoplasm (GO:0005654) | 1075 | 369 | 322.62 | 1.14 | 6.33E-05 | 6.92E-03 |
| ***PANTHER GO-Complete Biological Process*** | | | | | | |
| mitotic cell cycle process (GO:1903047) | 181 | 82 | 54.32 | 1.51 | 7.06E-06 | 2.03E-02 |
| cell cycle process (GO:0022402) | 267 | 118 | 80.13 | 1.47 | 2.23E-07 | 1.93E-03 |
| mitotic cell cycle (GO:0000278) | 210 | 92 | 63.02 | 1.46 | 9.87E-06 | 2.13E-02 |
| cell cycle (GO:0007049) | 372 | 154 | 111.64 | 1.38 | 4.77E-07 | 2.06E-03 |

**Table S6 – GO enrichment analysis results – proteins with phosphosites enriched in mitosis**
