## Supplementary Table S7 for "Spindle morphology changes between meiosis and mitosis driven by CK2 regulation of the Ran pathway"

| Gene ID | UNIPROT ID | Gene Name | Gene symbol |
|---|---|---|---|
| HUMAN\|HGNC=25309\|UniProtKB=Q6P2H3 | Q6P2H3 | Centrosomal protein of 85 kDa | CEP85 |
| HUMAN\|HGNC=25886\|UniProtKB=Q6NZ67 | Q6NZ67 | Mitotic-spindle organizing protein 2B | MZT2B |
| HUMAN\|HGNC=2584\|UniProtKB=Q9NQC7 | Q9NQC7 | Ubiquitin carboxyl-terminal hydrolase CYLD | CYLD |
| HUMAN\|HGNC=1722\|UniProtKB=P06493 | P06493 | Cyclin-dependent kinase 1 | CDK1 |
| HUMAN\|HGNC=27102\|UniProtKB=Q86Y91 | Q86Y91 | Kinesin-like protein KIF18B | KIF18B |
| HUMAN\|HGNC=20778\|UniProtKB=P07437 | P07437 | Tubulin beta chain | TUBB |
| HUMAN\|HGNC=24488\|UniProtKB=Q8NBT0 | Q8NBT0 | POC1 centriolar protein homolog A | POC1A |
| HUMAN\|HGNC=11354\|UniProtKB=Q8WVM7 | Q8WVM7 | Cohesin subunit SA-1 | STAG1 |
| HUMAN\|HGNC=21862\|UniProtKB=Q8WVS4 | Q8WVS4 | Cytoplasmic dynein 2 intermediate chain 1 | DYNC2I1 |
| HUMAN\|HGNC=29130\|UniProtKB=O94927 | O94927 | HAUS augmin-like complex subunit 5 | HAUS5 |
| HUMAN\|HGNC=20262\|UniProtKB=Q8IX90 | Q8IX90 | Spindle and kinetochore-associated protein 3 | SKA3 |
| HUMAN\|HGNC=24489\|UniProtKB=Q8ND56 | Q8ND56 | Protein LSM14 homolog A | LSM14A |
| HUMAN\|HGNC=1149\|UniProtKB=O60566 | O60566 | Mitotic checkpoint serine_threonine-protein kinase BUB1 beta | BUB1B |
| HUMAN\|HGNC=27000\|UniProtKB=Q8IVT2 | Q8IVT2 | Mitotic interactor and substrate of PLK1 | MISP |
| HUMAN\|HGNC=9341\|UniProtKB=O43663 | O43663 | Protein regulator of cytokinesis 1 | PRC1 |
| HUMAN\|HGNC=20772\|UniProtKB=Q13509 | Q13509 | Tubulin beta-3 chain | TUBB3 |
| HUMAN\|HGNC=30046\|UniProtKB=Q96BK5 | Q96BK5 | PIN2_TERF1-interacting telomerase inhibitor 1 | PINX1 |
| HUMAN\|HGNC=18745\|UniProtKB=Q9Y6G9 | Q9Y6G9 | Cytoplasmic dynein 1 light intermediate chain 1 | DYNC1LI1 |
| HUMAN\|HGNC=7723\|UniProtKB=Q8NHV4 | Q8NHV4 | Protein NEDD1 | NEDD1 |
| HUMAN\|HGNC=795\|UniProtKB=Q13315 | Q13315 | Serine-protein kinase ATM | ATM |
| HUMAN\|HGNC=13516\|UniProtKB=Q9NR09 | Q9NR09 | Baculoviral IAP repeat-containing protein 6 | BIRC6 |
| HUMAN\|HGNC=14074\|UniProtKB=Q9NZ56 | Q9NZ56 | Formin-2 | FMN2 |
| HUMAN\|HGNC=17088\|UniProtKB=Q7Z460 | Q7Z460 | CLIP-associating protein 1 | CLASP1 |
| HUMAN\|HGNC=16864\|UniProtKB=Q15398 | Q15398 | Disks large-associated protein 5 | DLGAP5 |
| HUMAN\|HGNC=3155\|UniProtKB=Q9H8V3 | Q9H8V3 | Protein ECT2 | ECT2 |
| HUMAN\|HGNC=29536\|UniProtKB=O60336 | O60336 | Mitogen-activated protein kinase-binding protein 1 | MAPKBP1 |
| HUMAN\|HGNC=6391\|UniProtKB=Q14807 | Q14807 | Kinesin-like protein KIF22 | KIF22 |
| HUMAN\|HGNC=30532\|UniProtKB=Q9BT25 | Q9BT25 | HAUS augmin-like complex subunit 8 | HAUS8 |
| HUMAN\|HGNC=30829\|UniProtKB=Q9BVA1 | Q9BVA1 | Tubulin beta-2B chain | TUBB2B |
| HUMAN\|HGNC=14629\|UniProtKB=Q53HL2 | Q53HL2 | Borealin | CDCA8 |
| HUMAN\|HGNC=17380\|UniProtKB=Q9UJX3 | Q9UJX3 | Anaphase-promoting complex subunit 7 | ANAPC7 |
| HUMAN\|HGNC=29022\|UniProtKB=Q69YQ0 | Q69YQ0 | Cytospin-A | SPECC1L |
| HUMAN\|HGNC=14540\|UniProtKB=Q9H4A3 | Q9H4A3 | Serine_threonine-protein kinase WNK1 | WNK1 |
| HUMAN\|HGNC=11524\|UniProtKB=Q9Y6A5 | Q9Y6A5 | Transforming acidic coiled-coil-containing protein 3 | TACC3 |
| HUMAN\|HGNC=2680\|UniProtKB=Q9BTC0 | Q9BTC0 | Death-inducer obliterator 1 | DIDO1 |
| HUMAN\|HGNC=6058\|UniProtKB=Q9NQS7 | Q9NQS7 | Inner centromere protein | INCENP |
| HUMAN\|HGNC=9077\|UniProtKB=P53350 | P53350 | Serine_threonine-protein kinase PLK1 | PLK1 |
| HUMAN\|HGNC=17619\|UniProtKB=Q9NXR1 | Q9NXR1 | Nuclear distribution protein nudE homolog 1 | NDE1 |
| HUMAN\|HGNC=1249\|UniProtKB=Q9ULW0 | Q9ULW0 | Targeting protein for Xklp2 | TPX2 |
| HUMAN\|HGNC=25948\|UniProtKB=Q7Z4H7 | Q7Z4H7 | HAUS augmin-like complex subunit 6 | HAUS6 |
| HUMAN\|HGNC=19422\|UniProtKB=O75528 | O75528 | Transcriptional adapter 3 | TADA3 |
| HUMAN\|HGNC=9804\|UniProtKB=Q9H0H5 | Q9H0H5 | Rac GTPase-activating protein 1 | RACGAP1 |
| HUMAN\|HGNC=1723\|UniProtKB=Q12834 | Q12834 | Cell division cycle protein 20 homolog | CDC20 |
| HUMAN\|HGNC=25083\|UniProtKB=Q8N0Z3 | Q8N0Z3 | Spindle and centriole-associated protein 1 | SPICE1 |
| HUMAN\|HGNC=7910\|UniProtKB=P06748 | P06748 | Nucleophosmin | NPM1 |
| HUMAN\|HGNC=1717\|UniProtKB=Q16181 | Q16181 | Septin-7 | SEPTIN7 |
| HUMAN\|HGNC=12401\|UniProtKB=P33981 | P33981 | Dual specificity protein kinase TTK | TTK |
| HUMAN\|HGNC=18538\|UniProtKB=Q9BXS6 | Q9BXS6 | Nucleolar and spindle-associated protein 1 | NUSAP1 |
| HUMAN\|HGNC=4201\|UniProtKB=Q92830 | Q92830 | Histone acetyltransferase KAT2A | KAT2A |
| HUMAN\|HGNC=9884\|UniProtKB=P06400 | P06400 | Retinoblastoma-associated protein | RB1 |
| HUMAN\|HGNC=15480\|UniProtKB=Q9NSV4 | Q9NSV4 | Protein diaphanous homolog 3 | DIAPH3 |
| HUMAN\|HGNC=1744\|UniProtKB=Q99741 | Q99741 | Cell division control protein 6 homolog | CDC6 |
| HUMAN\|HGNC=9787\|UniProtKB=O95235 | O95235 | Kinesin-like protein KIF20A | KIF20A |
| HUMAN\|HGNC=20771\|UniProtKB=P68371 | P68371 | Tubulin beta-4B chain | TUBB4B |
| HUMAN\|HGNC=28959\|UniProtKB=Q14008 | Q14008 | Cytoskeleton-associated protein 5 | CKAP5 |
| HUMAN\|HGNC=13339\|UniProtKB=O95239 | O95239 | Chromosome-associated kinesin KIF4A | KIF4A |
| HUMAN\|HGNC=6393\|UniProtKB=Q99661 | Q99661 | Kinesin-like protein KIF2C | KIF2C |
| HUMAN\|HGNC=4425\|UniProtKB=Q08379 | Q08379 | Golgin subfamily A member 2 | GOLGA2 |
| HUMAN\|HGNC=12759\|UniProtKB=Q9P2S5 | Q9P2S5 | WD repeat-containing protein WRAP73 | WRAP73 |
| HUMAN\|HGNC=1728\|UniProtKB=P30260 | P30260 | Cell division cycle protein 27 homolog | CDC27 |
| HUMAN\|HGNC=3338\|UniProtKB=Q14247 | Q14247 | Src substrate cortactin | CTTN |
| HUMAN\|HGNC=1990\|UniProtKB=Q8WWK9 | Q8WWK9 | Cytoskeleton-associated protein 2 | CKAP2 |
| HUMAN\|HGNC=17008\|UniProtKB=Q92547 | Q92547 | DNA topoisomerase 2-binding protein 1 | TOPBP1 |
| HUMAN\|HGNC=28109\|UniProtKB=Q96BD8 | Q96BD8 | Spindle and kinetochore-associated protein 1 | SKA1 |

**Table S7 – Proteins in Spindle GO category (GO:0005819) with phosphosites enriched in mitosis relative to meiosis**
