## Supplementary Table S8 for "Spindle morphology changes between meiosis and mitosis driven by CK2 regulation of the Ran pathway"

| Protein name (*Xenopus*) | Uniprot ID (Human) | Protein | Protein Name | Ran-MT Proteome (Rosas-Salvans et al., 2018) | HeLa Spindle Proteome (Rao et al., 2018) | MiCroKITs database (Huang et al., 2015) | Localization from literature search | Normalization |
|---|---|---|---|---|---|---|---|---|
| rna77834 | A0A0A0MSW3 | A0A0A0MSW3 | Nuclear pore complex protein Nup214 | | | | | BOTH |
| rna28203 | P23443 | KS6B1 | Ribosomal protein S6 kinase beta-1 | | | | | BOTH |
| rna57707 | Q9NVR7 | TBCC1 | TBCC domain-containing protein 1 | | | Centrosome | Centrosome; Spindle | BOTH |
| rna23924 | E9PGM1 | E9PGM1 | Eukaryotic translation initiation factor 4 gamma 1 | | + | | | BOTH |
| rna57206 | E9PGM1 | E9PGM1 | Eukaryotic translation initiation factor 4 gamma 1 | | + | | | BOTH |
| rna33897 | H7BZN3 | H7BZN3 | Telomere-associated protein RIF1 | | | | Spindle | BOTH |
| rna1597 | P23443 | KS6B1 | Ribosomal protein S6 kinase beta-1 | | | | | BOTH |
| rna22763 | Q9Y2D8 | ADIP | Afadin- and alpha-actinin-binding protein | + | + | Centrosome (predicted from orthologs) | Centrosome; Spindle | BOTH |
| rna57828 | H3BNU8 | H3BNU8 | Lysine-rich nucleolar protein 1 | | | | | BOTH |
| rna41108 | Q9Y2T7 | YBOX2 | Y-box-binding protein 2 | + | + | | | BOTH |
| rna3502 | Q06190 | P2R3A | Serine/threonine-protein phosphatase 2A regulatory subunit B\'\' subunit alpha | | | | | BOTH |
| rna39157 | Q96QE3 | ATAD5 | ATPase family AAA domain-containing protein 5 | | + | | Centrosome | BOTH |
| rna84769 | A0A6Q8PHA4 | A0A6Q8PHA4 | Transcriptional regulator ATRX | | | | | BOTH |
| rna2319 | O95049 | ZO3 | Tight junction protein ZO-3 | | | | | BOTH |
| rna18636 | D6RD46 | D6RD46 | LIM and calponin homology domains-containing protein 1 | | | | | BOTH |
| rna78702 | Q9UJF2 | NGAP | Ras GTPase-activating protein nGAP | + | + | | | BOTH |
| rna7007 | E9PPJ0 | E9PPJ0 | Splicing factor 3B subunit 2 | | + | | | PHASE ONLY |
| rna83354 | Q9BW71 | HIRP3 | HIRA-interacting protein 3 | | | | Spindle | BOTH |
| rna57446 | O75496 | GEMI | Geminin | | + | Centrosome | Centrosome | BOTH |
| rna85185 | Q9UKK3 | PARP4 | Protein mono-ADP-ribosyltransferase PARP4 | | | | Spindle | BOTH |
| rna50143 | Q9NYZ3 | GTSE1 | G2 and S phase-expressed protein 1 | + | + | | Microtubules | BOTH |
| rna30741 | Q9BW71 | HIRP3 | HIRA-interacting protein 3 | | | | Spindle | PHASE ONLY |
| rna78346 | O15541 | R113A | E3 ubiquitin-protein ligase RNF113A | | | | | BOTH |
| rna94082 | E9PCT1 | E9PCT1 | Serine/arginine repetitive matrix protein 1 | | + | | | BOTH |
| rna74416 | Q9BWT3 | PAPOG | Poly(A) polymerase gamma | | | | | BOTH |
| rna54475 | K7ES11 | K7ES11 | Ubiquitin conjugating enzyme E2 O | | | | | BOTH |
| rna35349 | B7ZM65 | B7ZM65 | PHD and RING finger domain-containing protein 1 | | | | | BOTH |
| rna29838 | Q9Y385 | UB2J1 | Ubiquitin-conjugating enzyme E2 J1 | | | | | BOTH |
| rna7284 | Q9UHB6 | LIMA1 | LIM domain and actin-binding protein 1 | + | + | Midbody | | BOTH |
| rna31688 | Q9NQS7 | INCE | Inner centromere protein | + | + | Midbody; Kinetochore; Spindle | Spindle; Kinetochore | BOTH |
| rna74412 | F2Z2T0 | F2Z2T0 | Rab-like protein 6 | | | | Centrosome | PHASE ONLY |
| rna33567 | B7Z683 | B7Z683 | Active breakpoint cluster region-related protein | | | | | PHASE ONLY |
| rna4231 | B7Z683 | B7Z683 | Active breakpoint cluster region-related protein | | | | | PHASE ONLY |
| rna30541 | A0A3B3IRV6 | A0A3B3IRV6 | Tight junction protein ZO-2 | | | | | PHASE ONLY |
| rna52899 | Q8WVC0 | LEO1 | RNA polymerase-associated protein LEO1 | | + | | Centrosome | PHASE ONLY |
| rna448 | I3L4D8 | I3L4D8 | Serine/arginine repetitive matrix protein 2 | | + | | | PHASE ONLY |
| rna20659 | F5H8D7 | F5H8D7 | DNA repair protein XRCC1 | | + | | Centrosome | PHASE ONLY |
| rna40394 | P18887 | XRCC1 | DNA repair protein XRCC1 | + | + | Centrosome | Centrosome | PHASE ONLY |
| rna85337 | Q9H7L9 | SDS3 | Sin3 histone deacetylase corepressor complex component SDS3 | | | | | BOTH |
| rna29250 | Q6VY07 | PACS1 | Phosphofurin acidic cluster sorting protein 1 | | | | | PHASE ONLY |
| rna25350 | Q13283 | G3BP1 | Ras GTPase-activating protein-binding protein 1 | | + | | Microtubules | PHASE ONLY |
| rna8756 | A0A3B3IS57 | A0A3B3IS57 | DNA helicase | | | | Centrosome | BOTH |
| rna87682 | Q9Y5S2 | MRCKB | Serine/threonine-protein kinase MRCK beta | | + | | | BOTH |
| rna35385 | O95235 | KI20A | Kinesin-like protein KIF20A | + | + | Midbody; Spindle | Spindle | BOTH |
| rna33120 | Q13547 | HDAC1 | Histone deacetylase 1 | + | + | Centrosome; Kinetochore | Centrosome; Kinetochore | PHASE ONLY |
| rna44245 | Q8NFH5 | NUP35 | Nucleoporin NUP35 | | + | | Spindle | PHASE ONLY |
| rna1994 | Q9Y2T7 | YBOX2 | Y-box-binding protein 2 | + | + | | | PHASE ONLY |
| rna23258 | Q13242 | SRSF9 | Serine/arginine-rich splicing factor 9 | | + | | | PHASE ONLY |
| rna30403 | Q9BWT1 | CDCA7 | Cell division cycle-associated protein 7 | | | | | BOTH |
| rna9276 | E9PCY5 | E9PCY5 | DNA topoisomerase 2 | | | | | PHASE ONLY |
| rna38444 | Q9Y2U8 | MAN1 | Inner nuclear membrane protein Man1 | | + | | | BOTH |
| rna929 | P24864 | CCNE1 | G1/S-specific cyclin-E1 | | | | Centrosome | PHASE ONLY |
| rna27729 | Q99741 | CDC6 | Cell division control protein 6 homolog | + | | | Centrosome | BOTH |
| rna10397 | Q96ST2 | IWS1 | Protein IWS1 homolog | | + | | | PHASE ONLY |
| rna33962 | A0A2R8Y4J3 | A0A2R8Y4J3 | AP-3 complex subunit delta | | | | | BOTH |
| rna12135 | Q6UN15 | FIP1 | Pre-mRNA 3\'-end-processing factor FIP1 | + | + | | | PHASE ONLY |
| rna55791 | K7EPA2 | K7EPA2 | Centrosomal protein of 192 kDa | | | | Centrosome | PHASE ONLY |
| rna53758 | Q9NPQ8 | RIC8A | Synembryn-A | | + | Centrosome; Midbody | Spindle; Centrosome | PHASE ONLY |
| rna36772 | Q86V48 | LUZP1 | Leucine zipper protein 1 | + | + | Centrosome (predicted from orthologs) | Centrosome; Spindle | BOTH |
| rna47394 | H0YCE8 | H0YCE8 | RNA polymerase-associated protein CTR9 homolog | | | | | BOTH |
| rna92132 | Q9NQS7 | INCE | Inner centromere protein | + | + | Midbody; Kinetochore; Spindle | Spindle; Kinetochore | PHASE ONLY |
| rna32873 | P36915 | GNL1 | Guanine nucleotide-binding protein-like 1 | | | | | PHASE ONLY |
| rna27096 | Q96T17 | MA7D2 | MAP7 domain-containing protein 2 | | | | Microtubules; Centrosome | PHASE ONLY |
| rna9600 | P05455 | LA | Lupus La protein | | | | | BOTH |
| rna49008 | P05455 | LA | Lupus La protein | | | | | BOTH |
| rna27700 | A0A087WUT6 | A0A087WUT6 | Eukaryotic translation initiation factor 5B | | | | | PHASE ONLY |
| rna41041 | O43818 | U3IP2 | U3 small nucleolar RNA-interacting protein 2 | | + | | | PHASE ONLY |
| rna7305 | E5RFQ8 | E5RFQ8 | Nucleoplasmin-2 | | | | | PHASE ONLY |
| rna18434 | E5RFQ8 | E5RFQ8 | Nucleoplasmin-2 | | | | | PHASE ONLY |
| rna4284 | Q5VT52 | RPRD2 | Regulation of nuclear pre-mRNA domain-containing protein 2 | | + | | | PHASE ONLY |
| rna89432 | Q5VT52 | RPRD2 | Regulation of nuclear pre-mRNA domain-containing protein 2 | | + | | | PHASE ONLY |
| rna39162 | Q86TB9 | PATL1 | Protein PAT1 homolog 1 | + | + | | | PHASE ONLY |
| rna89349 | Q53H80 | AKIR2 | Akirin-2 | | | | | PHASE ONLY |
| rna90230 | Q53H80 | AKIR2 | Akirin-2 | | | | | PHASE ONLY |
| rna42755 | M0R0Q7 | M0R0Q7 | DNA ligase | | + | | | PHASE ONLY |
| rna35230 | Q7RTY7 | OVCH1 | Ovochymase-1 | | | | | TOTAL ONLY |
| rna19173 | Q9NRA8 | 4ET | Eukaryotic translation initiation factor 4E transporter | + | | | | TOTAL ONLY |
| rna53087 | Q9H211 | CDT1 | DNA replication factor Cdt1 | | + | Kinetochore | Kinetochore | TOTAL ONLY |

### Table S8 – Candidate proteins annotated with localization data
